# Promoters constrain evolution of expression levels of essential genes in *Escherichia coli*

**DOI:** 10.1101/2024.05.20.594948

**Authors:** Saburo Tsuru, Naoki Hatanaka, Chikara Furusawa

**Affiliations:** Universal Biology Institute, Graduate School of Science, The University of Tokyo, 7-3-1 Hongo, Bunkyo-ku, Tokyo 113-0033, Japan; Department of Physics, Graduate School of Science, The University of Tokyo, 7-3-1 Hongo, Bunkyo-ku, Tokyo 113-0033, Japan; Center for Biosystems Dynamics Research (BDR), RIKEN, 6-2-3 Furuedai, Suita, Osaka 565- 0874, Japan

## Abstract

Variability in expression levels in response to random genomic mutations varies among genes, influencing both the facilitation and constraint of phenotypic evolution in organisms. Despite its importance, both the underlying mechanisms and evolutionary origins of this variability remain largely unknown due to the mixed contributions of *cis*- and *trans*-acting elements. To address this issue, we focused on the mutational variability of *cis*-acting elements, that is, promoter regions, in *Escherichia coli*. Random mutations were introduced into the natural and synthetic promoters to generate mutant promoter libraries. By comparing the variance in promoter activity of these mutant libraries, we found no significant difference in mutational variability in promoter activity between promoter groups, suggesting the absence of a signature of natural selection for mutational robustness. In contrast, the promoters controlling essential genes exhibited a remarkable bias in mutational variability, with mutants displaying higher activities than the wild types being relatively rare compared to those with lower activities. Our evolutionary simulation on a rugged fitness landscape provided a rationale for this vulnerability. These findings suggest that past selection created non-uniform mutational variability in promoters biased toward lower activities of random mutants now constrain future evolution of downstream essential genes toward higher expression levels.

## Introduction

The rate and direction of phenotypic evolution largely depend on phenotypic variability, the tendency to vary phenotypically (Lynch and Walsh 1998; Walker 2007). It is widely recognized that organisms often exhibit nonuniform phenotypic variability, with certain variants arising more frequently than others in response to different genetic and environmental perturbations, even in the absence of natural selection(Darwin 1859; Waddington 1957; Smith, et al. 1985; Noble, et al. 2019). Therefore, it is crucial to elucidate the mechanisms underlying the bias in phenotypic variability to understand why phenotypic evolution proceeds as it does(Uller, et al. 2018). Transcriptional variability of genes across different perturbations plays a significant role in phenotypic evolution because the evolution of many traits in organisms largely depends on the evolution of expression levels and their regulation(Wittkopp, et al. 2004; Zheng, et al. 2011). Interestingly, previous studies have identified differences in transcriptional variability from gene to gene in response to various perturbations(Denver, et al. 2005; Rifkin, et al. 2005; Landry, et al. 2007; Tsuru and Furusawa 2024), where genes related to cellular growth and maintenance tend to exhibit lower transcriptional variability against random genetic perturbations to the genome in yeast(Landry, et al. 2007) and bacteria(Tsuru and Furusawa 2024). These studies, supported by theoretical evidence(Kaneko 2009; Draghi and Whitlock 2012; Furusawa and Kaneko 2018), suggest that the transcriptional variability of genes is not a random occurrence devoid of biological significance but rather a product of natural selection.

What molecular mechanisms account for gene-to-gene differences in transcriptional variability? In principle, both *cis*- and *trans*-acting elements can influence transcriptional variability(Wittkopp, et al. 2004; Tirosh, et al. 2006; Landry, et al. 2007; Payne and Wagner 2015; Tsuru and Furusawa 2024). Previous studies(Denver, et al. 2005; Rifkin, et al. 2005; Landry, et al. 2007; Tsuru and Furusawa 2024) explored the transcriptional variability against different genetic perturbations using mutation accumulation (MA) experiments(Halligan and Keightley 2009), where independent mutants were generated by accumulating genomic mutations through genetic drift, and their transcriptome profiles were obtained under identical environmental conditions. Although MA experiments provide a powerful method to quantify transcriptional variability in the absence of strong natural selection, the indiscriminate accumulation of mutations on *cis*- and *trans*-acting elements(Tsuru, et al. 2015) complicates the identification of the molecular mechanism underlying transcriptional variability. Accordingly, to overcome this difficulty, it is necessary to consider the contributions of *cis*- and *trans*-acting elements separately. Although such approaches have been extensively employed in yeast(Hornung, et al. 2012; Metzger, et al. 2015; Duveau, et al. 2021), it remains largely unknown which elements practically contribute to gene-to-gene differences in transcriptional variability associated with functions of gene products.

Why have genes related to cellular growth and maintenance acquired low mutational variability? The evolutionary origin of mutational variability is debatable (de Visser, et al. 2003). Mutational variability could result from direct natural selection for mutational robustness(Waddington 1942), or could be a byproduct of selection for robustness against environmental perturbations(Meiklejohn and Hartl 2002; de Visser, et al. 2003). Several biological traits, such as metabolic flux(Ho and Zhang 2016) and cellular morphology(Ho and Zhang 2014), show signs of adaptive evolution through direct selection for mutational robustness, where traits highly related to cellular growth fitness are likely to acquire lower mutational variability through direct selection for mutational robustness. On the other hand, other studies have identified congruence in trait variability between environmental and genetic perturbations, such as in RNA structure(Szollosi and Derenyi 2009) and gene expression levels(Landry, et al. 2007; Tsuru and Furusawa 2024), supporting the relevance of the byproduct of selection for environmental robustness. As an alternative scenario, mutational variability may spontaneously emerge with trait evolution, regardless of selection for robustness(Siegal and Bergman 2002; de Visser, et al. 2003). To date, the most appropriate scenario for low transcriptional variability of genes related to cellular growth and maintenance remains largely unknown.

To address these questions, we focused on the mutational variability governed by cis-acting elements, that is, promoter regions, of genes in *Escherichia coli*. We created a mutant library for each promoter through random mutagenesis and analyzed the changes in promoter activity from the wild type to mutant using flow cytometry. To explore the relevant evolutionary scenarios for the mutational variability of promoters, we compared promoters controlling essential genes for cellular growth with those controlling nonessential genes. These groups were expected to experience different natural selection pressures on their transcriptional variability in the past. In addition, random sequences were used to construct synthetic promoters that experienced no selection for mutational robustness. Interestingly, when comparing the variance in promoter activity of the mutant libraries, these three promoter groups exhibited similar mutational variability in promoter activity, indicating the absence of signatures of natural selection for mutational robustness. In contrast, the promoters for essential genes showed a remarkable vulnerability to mutations compared to the other promoter groups, where variants with higher activity than the wild type arose less frequently than variants with lower activity. Our evolutionary simulation on a rugged fitness landscape explains the higher vulnerability of the promoters for essential genes. These results suggest that past selection created a bias in mutational variability in promoters now constrains future evolution of downstream essential genes toward higher expression levels.

## Results

### Higher expression levels and lower transcriptional variability of essential genes in *E. coli*

First, we analyzed the transcriptional variability of essential genes in *E. coli* in response to various genomic mutations (**Fig. 1A**). Previously(Tsuru and Furusawa 2024), we obtained transcriptome profiles of six independent mutants that had accumulated genomic mutations through genetic drift via the mutation accumulation (MA) experiment(Tsuru, et al. 2015), termed the Mut dataset. Using this dataset, we quantified the mean and standard deviation of the expression levels for each of thousands of genes (**Fig. 1B**). To compensate for the nonmonotonic mean-dependency of the standard deviations, we calculated the vertical distance from the smoothed spline of the running median of the standard deviations, termed DM_mut_. The DM_mut_ values reflect the mean transcriptional variability in response to different genetic perturbations(Tsuru and Furusawa 2024). We categorized thousands of genes into essential and nonessential genes for cell growth, following the definition of gene essentiality by Goodall *et al*(Goodall, et al. 2018). We confirmed that essential genes showed higher expression levels than nonessential genes across the different mutants (**Fig. 1C**). We also found that essential genes had lower DM_mut_ than nonessential genes (**Fig. 1D**).

**Fig. 1:**
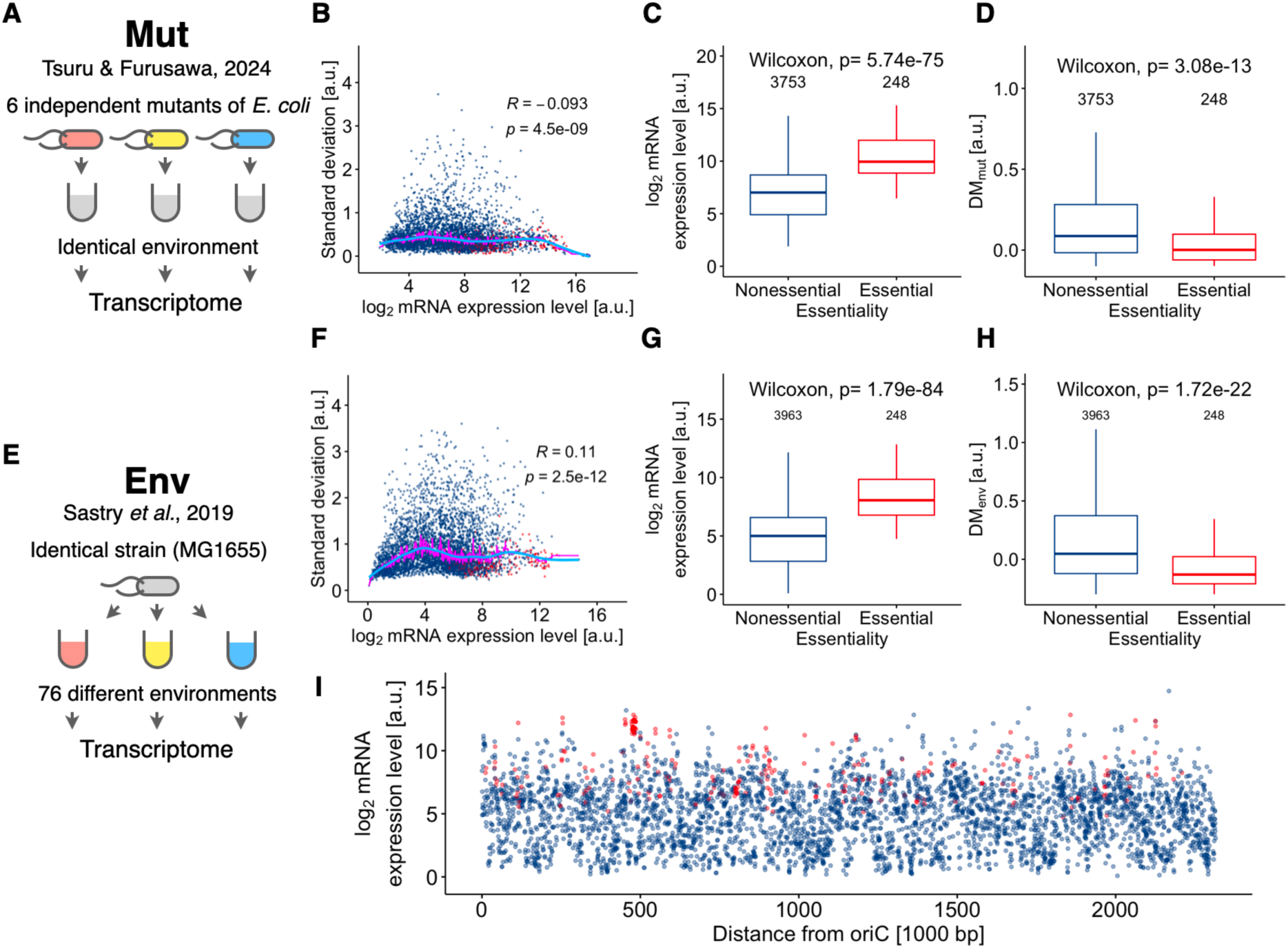
Transcriptional variability of essential genes in *E. coli*. (**A**–**D**) Transcriptional profiles of six independent mutants of *E. coli* cultured under identical environmental conditions, termed the Mut dataset. The mutants were created through the MA experiment to accumulate random genomic mutations by genetic drift. (**E**–**I**) Transcriptional profiles of a single strain, MG1655, cultured under 76 different environmental conditions, termed the Env dataset. (**A**, **E**) Schematics of the datasets. (**B**, **F**) Relationship between the means and standard deviations of mRNA expression levels for thousands of genes. The magenta and cyan lines represent running medians and their smoothed splines, respectively. Spearman’s rank correlation coefficients (R) and p-values are shown. (**C**, **G**) Mean expression levels of essential and nonessential genes. (**D**, **H**) Transcriptional variability, represented as either DM_mut_ or DM_env_, of essential and nonessential genes. The DM values were calculated as the vertical distance of the standard deviations from the smoothed splines of the running median of standard deviations in panels **B** and **F**. The lower and upper edges of the boxes in panels **C**, **D**, **G**, **H** represent the first (q1) and third (q3) quartiles, respectively. The horizontal lines in the boxes represent the medians (m). The whiskers from the boxes extend to the most extreme observed values inside inner fences, m±1.5(q3–q1). (**I**) Relationship between mean expression levels and the shorter distance from *oriC* on the *E. coli* chromosome. Red and blue dots in panels **B**, **F**, and **I** represent essential and nonessential genes, respectively.

Next, we explored the transcriptional variability of essential genes in response to different environmental perturbations (**Fig. 1E,F**) using a public dataset(Sastry, et al. 2019; Rychel, et al. 2021) of the transcriptome profiles of a single strain of *E. coli*, MG1655, cultured under 76 different environmental conditions, termed the Env dataset. Previously(Tsuru and Furusawa 2024), we quantified the mean-controlled standard deviations, termed DM_env_, using a method similar to DM_mut_. The DM_env_ reflects transcriptional variability in response to different environmental perturbations. We confirmed that the essential genes exhibited higher expression levels (**Fig. 1G**) and lower variability (**Fig. 1H**) than the nonessential genes across different environmental conditions. Taken together, essential genes exhibited higher expression levels and lower transcriptional variability in response to different perturbations, which is similar to the characteristics of essential genes in yeast(Tirosh, et al. 2006).

In *E. coli*, genes with many chromosomal copies or genes located at near the origin of replication (*oriC*) tend to have higher expression levels(Ying, et al. 2014). However, essential genes shown in **Fig. 1** are present as only single copies on the chromosome. In addition, these essential genes are widely spread on the chromosome and exhibited higher expression levels than neighboring nonessential genes over the chromosome (**Fig. 1I**). Therefore, gene dosage did not seem to explain the higher expression levels of essential genes. To explore whether the higher expression levels of essential genes are caused by chromosomal position or by higher-order chromosomal structures, we analyzed promoter-mediated gene expression from a plasmid based on the experimental dataset obtained by Silander *et al*(Silander, et al. 2012). Using flow cytometry, Silander *et al* quantified promoter activity of plasmid copies of different promoter regions in *E. coli*, where green fluorescent protein (GFP) was transcriptionally fused to plasmid copies of promoters. Based on fluorescence intensity from GFP, we compared promoter activity between promoters controlling essential genes (termed essential promoters) and those controlling nonessential genes only (termed nonessential promoters) (**supplementary fig. S1A**). We found that essential promoters tended to have higher activities than nonessential promoters in spite of plasmid copies. This result highlighted the importance of promoter regions in accounting for higher expression levels of essential genes independent of chromosomal positions or structures.

Like eukaryotes, bacteria often employ postreplicative DNA methylation for the epigenetic control of gene expression. In *E. coli*, DNA adenine methyltransferase plays a major role in the epigenetic control and the target GATC motifs are enriched in promoter regions(Oshima, et al. 2002). Despite that adenine methylation at GATC sites in promoters can inhibit transcription for some genes(Casadesus and Low 2006), we found that differences in promoter activity between essential and nonessential promoters were almost independent of the presence or absence of GATC motifs in the promoter regions (**supplementary fig. S1B**). These results implied that the higher expression levels of essential genes reflected higher promoter activities of essential promoters without known epigenetic controls.

### Construction of mutant libraries of *E. coli* promoters to measure mutational effects

To explore the molecular mechanisms and signs of natural selection underlying the lower transcriptional variability of essential genes, we focused on the promoter regions of genes with different essentialities. In particular, we explored how promoter activity varies in response to new mutations in the promoter regions and how this variability relates to essentiality of the downstream genes. To this end, we used 56 natural promoters from the *E. coli* Promoter Collection(Zaslaver, et al. 2006). Importantly, to ensure an unbiased comparison of changes in promoter activities between different essentialities and to avoid measurement of promoter activity below the lower detection limit in flow cytometry, we avoided the use of natural promoters with very low activity that were slightly enriched in the promoters of nonessential genes. To achieve this, we selected natural promoters with moderate to high activity before mutagenesis. This selection was based on the known promoter activities of the collection as measured by Silander et al(Silander, et al. 2012). Using the RegulonDB database(Tierrafria, et al. 2022), we retrieved information regarding known transcriptional units regulated by these promoters. Based on the presence of essential genes in the transcription units, we categorized natural promoters into nonessential and essential promoters, termed Nes and Ess, respectively (**Fig. 2A**). Nonessential promoters control transcriptional units comprising only nonessential genes, whereas essential promoters regulate those that include at least a single essential gene. In addition to these natural promoters, we created synthetic promoters, termed Syn, as controls that experienced no or minimal selection for mutational robustness. To this end, we placed random sequences (∼200 bp) upstream of the *gfp* gene on a low-copy-number plasmid, pUA66- mCherry (**Fig. 2B**, **supplementary fig. S2**). The library of mixed transformants containing this plasmid was analyzed using flow cytometry. The top 7% of cells showing higher expression levels of GFP (red shaded area in **Fig. 2B**) were sorted, and 16 unique clones were subsequently isolated. Importantly, these synthetic promoters experienced selection for their expression levels only once, and no longer experienced further mutagenesis or selection for expression level. Accordingly, they can be regarded as a control group that is almost free from selection for mutational robustness. Consequently, in the upstream of the GFP gene on the plasmid, we constructed three promoter groups, Nes, Ess, and Syn, comprising different sequences with low homology (**supplementary fig. S3, supplementary note**). The green fluorescence from GFP was used as a measure of promoter activity(Zaslaver, et al. 2006; Silander, et al. 2012). Despite the diversity in sequence, we confirmed that the mean activities of the three promoter groups were comparable to each other, as designed (**Fig. 2C**). We also confirmed that the obtained promoter activities were highly consistent with those measured by Silander et al(Silander, et al. 2012) (Spearman’s R=0.94, p<0.05, **supplementary fig. S4A**). Compared to the Env dataset, the activities of the natural promoters were positively correlated with known mRNA expression levels of the genes downstream of the chromosomal promoters in the wild-type strain (Spearman’s R=0.64, **supplementary fig. S4B**), supporting that the wild-type promoter activities monitored by our method considerably reflect the activities of chromosomal promoters and their impact on the expression levels of downstream genes.

**Fig. 2:**
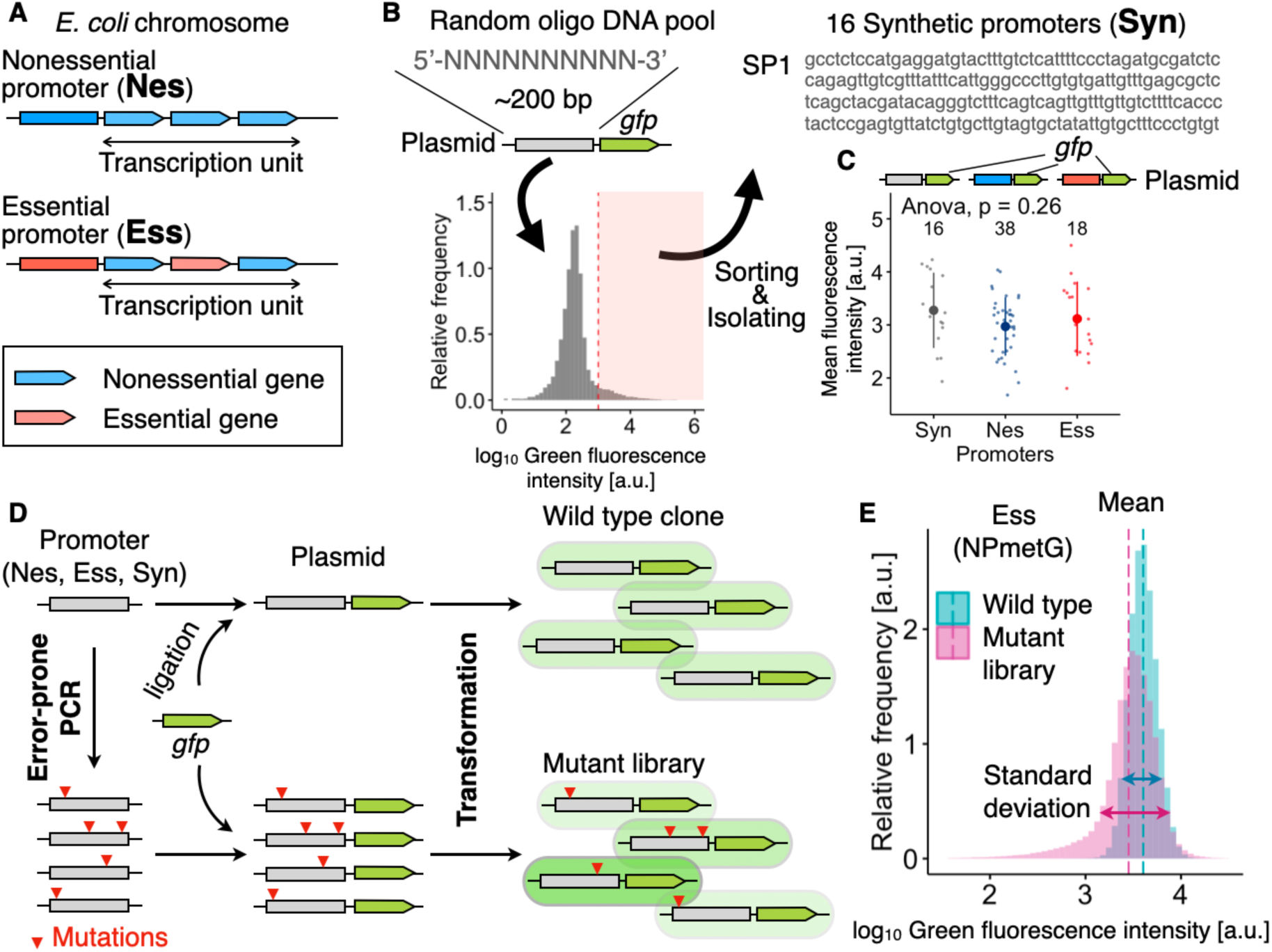
Experimental design to measure expression levels of mutant promoter library. (**A**) Natural promoters of *E. coli* were categorized into nonessential and essential promoters, termed Nes and Ess, respectively. Nonessential promoters control transcriptional units comprising only nonessential genes, while essential promoters regulate transcriptional units containing at least one essential gene. Gene essentiality was based on the requirement for cell growth, as defined by Goodall et al(Goodall, et al. 2018). The promoter regions were obtained from the *E. coli* Promoter Collection(Zaslaver, et al. 2006). (**B**) Synthetic promoters, termed Syn, were derived from a random oligo DNA pool. Transformants harboring a plasmid on which the random sequences were placed upstream of the *gfp* gene were analyzed using flow cytometry. The 16 unique transformants, satisfying a defined green fluorescence (red shaded area), were isolated. The obtained synthetic promoters were named SP1–16. (**C**) Natural and synthetic promoters were placed upstream of *gfp* on a plasmid. Transformants harboring each of these plasmids were analyzed using flow cytometry. Mean green fluorescence of each population is shown as small dots. The large dots and error bars represent the means and standard deviations within the groups, respectively. The number of promoters is indicated inside the panel. The p-value corresponds to ANOVA. (**D**) Schematic showing construction of mutant library from a given promoter region. The promoter region was subjected to random mutagenesis by error-prone PCR. A mutant library was created through transformation to yield more than 100,000 genetically-different mutants in a population. Unmutagenized plasmid was also transformed to create wild type clones. (**E**) Representative examples of log_10_-transformed green fluorescence intensity distributions of wild type and mutant library for the promoter region of the *metG* gene in *E. coli* (NPmetG). The means (vertical lines) and standard deviations (horizontal arrows) were calculated in each population.

For each of these promoters, we introduced mutations by error-prone PCR at a low frequency (6.3 mutations/kb on average, **Fig. 2D**) and integrated them into pUA66-mCherry to create a mutant library. Each mutant library had a genetic diversity of 10^5^ or more genotypes. The promoter activity of each mutant library was measured by flow cytometry based on the green fluorescence derived from downstream *gfp*, and was compared with the green fluorescence of the corresponding wild type (**Fig. 2E**). The introduction of mutations was limited to the promoter regions of the plasmid, while the chromosomal copies of the natural promoters were intact to avoid any confounding effects. The green fluorescence intensity was log_10_-transformed after subtracting the background (**supplementary fig. S5**), and the mean and standard deviation of the GFP distribution were calculated (**Fig. 2E, supplementary fig. S6**).

### Mutational variability in promoter activity was similar between different promoter groups

We confirmed that all mutant libraries showed larger variations than the corresponding wild-type clones, as designed (**Fig. 3A**), supporting that the difference in variations of GFP distributions between wild types and the mutant libraries reflected new genetic variations introduced by error-prone PCR rather than environmental changes or cell-to-cell heterogeneity. In addition, we also confirmed the well-known negative correlation between mean expression levels and standard deviations in GFP distributions regardless of the introduction of mutations or whether the promoters were natural or synthetic (**Fig. 3B,C**), which is consistent with previous observations(Zaslaver, et al. 2006; Wolf, et al. 2015). To compare different promoters, we compensated for the mean-dependency of the standard deviations of both wild types and mutant libraries by subtracting the regression line obtained from wild types (**Fig. 3B,C**). We subsequently calculated MV_prm_, a measure of mutational variability in promoter activity, by subtracting the compensated standard deviation of wild types from that of mutant libraries. Unexpectedly, MV_prm_, showed similar values across the three promoter groups (Wilcoxon test, adjusted p-values>0.05), even though each promoter might show unique values (**Fig. 3D**). These results indicated that mutational variability of the focal essential promoters showed little detectable signature of natural selection for mutational robustness despite their importance in cellular growth, even though such selection might be active. Moreover, MV_prm_ showed no significant positive correlations with DM_env_ and DM_mut_ of the downstream genes (**Fig. 3E,F**). The DM values were defined in **Fig. 1**. Using the RegulonDB and EcoCyc(Keseler, et al. 2021) databases to retrieve the known direct regulatory relationships between genes, we counted the number of direct and unique regulations that each promoter receives from transcriptional regulators and explored the impact of the number of regulations on mutational variability. We found that a larger number of activatory transcriptional regulators facilitated mutational variability in promoter activity (Spearman’s R=0.31, **Fig. 3G**), probably due to a larger number of mutational target sites. The relationship between MV_prm_ and the number of inhibitory transcriptional regulators was unclear because promoters with null regulation dominated the results (**Fig. 3H**). Despite that the number of activations was likely to make a difference in MV_prm_, the lack of significant correlations between MV_prm_ and DM values suggests that the mutational variability of promoters provides no considerable explanation for the observed gene- to-gene differences in transcriptional variability of the downstream genes in response to different perturbations at the genome scale (**Fig. 1**). These results suggest that the gene-to-gene differences in transcriptional variability in response to genome-wide perturbations may reflect a certain *trans*-acting molecular mechanism, mainly other than the *cis*-acting one, consistent with previous observations(Tsuru and Furusawa 2024). This implication was also rational because promoter regions are shorter than trans-acting mutational sites for a single gene.

**Fig. 3:**
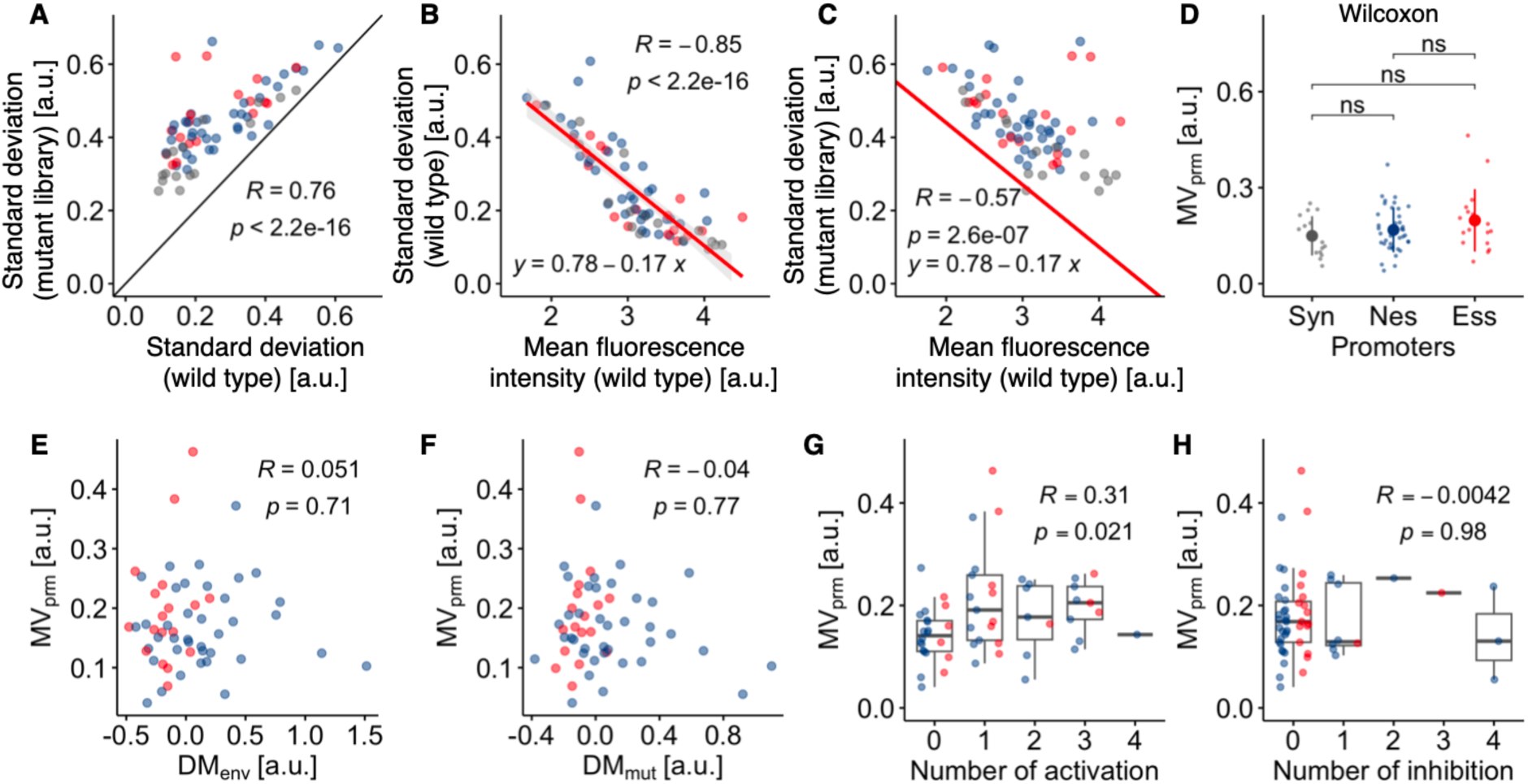
Mutational variability in promoter activity. (**A**) Comparison of the standard deviations in the GFP distributions between wild types and mutant libraries. The solid line represents y=x. (**B**, **C**) Relationship between the mean and standard deviation of the GFP distributions for wild types (**B**) and mutant libraries (**C**). Red lines represent a regression line for wild types and were used to compensate for the mean- dependency of the standard deviations of both wild types and mutant libraries by subtraction. (**D**) Mutational variability in promoter activity, term MV_prm_, is shown for each promoter group. The MV_prm_ was calculated by subtracting the compensated standard deviations of wild types from those of mutant libraries. The large dots and error bars are the means and standard deviations, respectively. Pairwise Wilcoxon test was examined, where ns represents no significance. (**E**, **F**) The relationships between MV_prm_ and the DM values (DM_env_ for **E** and DM_mut_ for **F**). The DM values were defined in Fig. 1. (**G**, **H**) Relationship between MV_prm_ and the number of known activation (**G**) and inhibition (**H**) of natural promoters by transcriptional regulators. The relationship between promoters and known transcriptional regulators were based on RegulonDB(Tierrafria, et al. 2022) and EcoCyc(Keseler, et al. 2021). The grey, blue, and red dots in each panel represent synthetic (Syn), nonessential (Nes), and essential (Ess) promoters (defined in Fig. 2), respectively. Spearman’s R and p-values are shown in panes **A**– **C** and **E**–**H**.

### Vulnerability in essential promotors against mutations

Next, we explored the directionality of the mutational changes in promoter activity. Consistent with the low mutation rate in error-prone PCR, the mean expression levels of mutant libraries were similar to those of wild types (Spearman’s R=0.99) (**Fig. 4A**). Nevertheless, we found that log_10_-fold changes in mean expression levels from wild types to mutant libraries followed a global tendency over expression levels, where the promoters with higher activities in wild types tended to decrease their activity in response to mutations (Spearman’s R=–056) (**Fig. 4B**). This tendency was retained even when the mean expression levels were calculated for the cells within the narrow gates with the fixed size over expression levels (**supplementary figs. S6,7**), supporting that the negative correlation was not a technical biproduct of our limitation of detecting a decrease in expression levels. This tendency may reflect bias in the mode, activation or inhibition, of transcriptional regulation(Kinney, et al. 2010; Belliveau, et al. 2018; Ireland, et al. 2020). We hypothesized that promoters that receive more activation than inhibition might tend to reduce their activity in response to mutations. To test this possibility, we defined activation bias as the number of known activations subtracted by the number of inhibitions (**supplementary table S1**). We found that the activation bias explained the observed mutational changes (**Fig. 4C**). Furthermore, the activation bias also explained the log_10_-fold changes after controlling the global trend using the regression line (**Fig. 4D**), indicating that a promoter becomes more vulnerable as it receives more activation among promoters with similar activities.

**Fig. 4:**
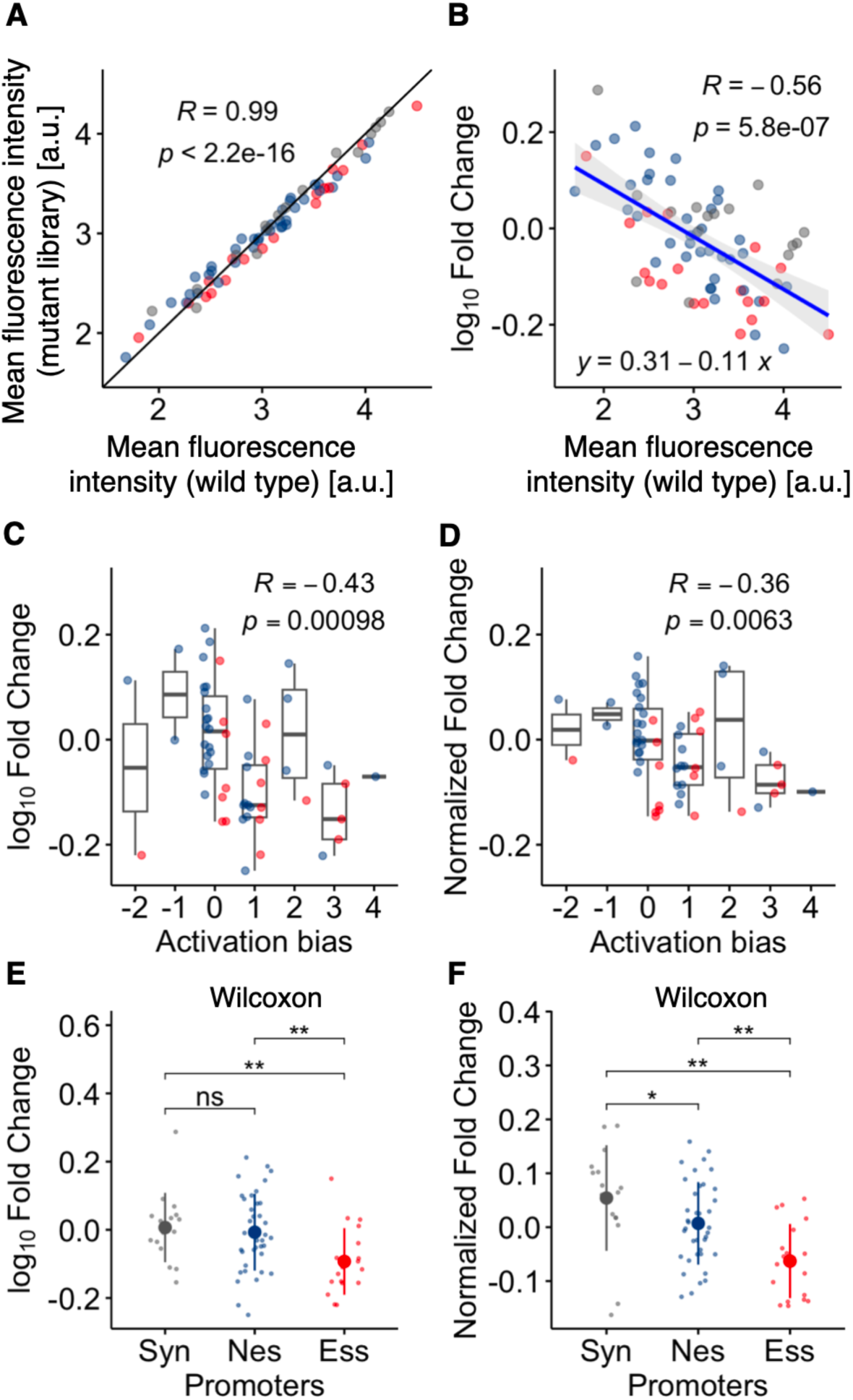
Biased mutational changes in activity of essential promoters. (**A**) Comparison of the means in the GFP distributions between wild types and mutant libraries. The solid line represents y=x. (**B**) Relationship between log_10_-fold change of mean expression levels from wild types to mutant libraries and the expression levels of wild types. The blue line represents a regression line used to compensate for the mean-dependency of the log_10_-fold changes by subtraction. (**C**, **D**) Relationship between activation bias in natural promoters and the unnormalized (**C**) and normalized (**D**) log_10_-fold changes. The activation bias was calculated by subtracting the number of known inhibitions from those of activations by transcriptional regulators. Spearman’s R and p-values are shown in panes **A–D**. Grey, blue, and red dots in each panel represent synthetic (Syn), nonessential (Nes), and essential (Ess) promoters (defined in Fig. 2), respectively. (**E**, **F**) Unnormalized (**E**) and normalized (**F**) log_10_-fold changes for each promoter group. The large dots and error bars are the means and standard deviations, respectively. Asterisks represent significance levels in pairwise Wilcoxon test (ns: p > 0.05; *: p ≤ 0.05; **: p ≤ 0.01). The p-values were adjusted using the BH method(Benjamini and Hochberg 1995).

Interestingly, synthetic and nonessential promoters showed similar mutational changes around zero, whereas essential promoters showed biased directionality in mutational changes toward negative values (**Fig. 4E**), even though the mean activities were almost the same among the three promoter groups (**Fig. 2C**). There were no significant differences in promoter length and GC content between the essential and nonessential promoters (Wilcoxon test, p>0.05). In addition, there was no significant difference in the mutation rate in error-prone PCR between the essential and nonessential promoters (**supplementary fig. S8A, supplementary table S2**). The mutational spectrum in error-prone PCR was quite similar between the two promoter groups (**supplementary fig. S8B**). These results supported the notion that the observed bias was not derived from a technical bias. This bias in essential promoters was retained for the normalized log_10_-fold changes (**Fig. 4F**). Accordingly, essential promoters were more vulnerable than other groups in terms of promoter activity. In other words, this biased directionality indicated the low frequency of the variants with higher activities than the wild-types in essential promoters. This biased variability might limit the transcriptional variability of essential genes in response to mutations in cis-regulatory regions, independently from transcriptional variability in response to genomic mutations (**Fig. 1**, **3F**).

### Vulnerability in essential promotors against mutations may reflect past selection for higher activity

The difference in mutational vulnerability between essential and nonessential promoters (**Fig. 4E, F**) could reflect the difference in past selection pressures acting on higher expression levels between essential and nonessential genes. Based on the observation that essential genes tended to have higher expression levels than nonessential genes (**Fig. 1**), we hypothesized that the remarkable vulnerability of essential promoters reflects local optima in terms of promoter activity. That is, essential promoters might have evolved toward higher activity through past evolution and might have been stuck in local optima because of the high essentiality of downstream genes. In contrast, nonessential promoters might have experienced relatively relaxed selection for higher expression levels because of the low essentiality of downstream genes. To explore the relevance of our assumption, we compared the impact of knockdown on growth fitness of *E. coli* between essential and nonessential genes using a public experimental dataset reported by Hawkins *et al*(Hawkins, et al. 2020). We found that knockdown of essential genes tended to be more deleterious for growth fitness than nonessential genes (**supplementary fig. S9A**), supporting our assumption that essential genes are under stronger selection pressure toward higher expression levels than nonessential genes. To investigate the emergence of differences in vulnerability depending on selection pressure, we performed a numerical simulation of promoter evolution on a simple rugged fitness landscape (**Fig. 5**). For simplicity, we modeled promoter genotype as 10-bit binary numbers such as 1101011011. That is, our model promoters are comprising of biallelic sites (0 or 1), which is simplified from real promoters comprising of quadallelic sites (A, T, G, and C). We assumed that the expression landscape is rugged and genotypes with higher expression levels are relatively rare, consistent with the previous experimental observations(Wolf, et al. 2015; Yona, et al. 2018). To satisfy this assumption, the mapping form genotype to expression level, equivalent to promoter activity, was defined by the Rough Mt. Fuji (RMF) model, as illustrated in **Fig. 5A, D**. In short, the genotype with the highest expression level, *g*^∗^, was first chosen at random. The expression levels of the other genotypes were decided as a decreasing linear function of the Hamming distance from *g*^∗^. Subsequently, stochastic noise was added to generate ruggedness. Finally, expression levels were set to range from 0 (lower bound) to 1 (upper bound) as detailed in the **Materials and Methods** section. Using this expression landscape, mutational changes were calculated for each genotype as the mean difference in the expression levels of all nearest neighbors with a single bit difference, or one Hamming distance, from the expression level of the focal genotype (**Fig. 5B, E**). This simple expression landscape captured the negative correlation between expression levels and mutational changes (**Fig. 5E**), which is consistent with the global trend observed in our experiment (**Fig. 4B**). This trend was likely due to the prevalence of local sinks (green dots) at lower expression levels and frequent local peaks (red dots) at higher expression levels (**Fig. 5B, E**). To clarify this point, we calculated the fraction of the nearest neighboring genotypes with higher expression levels for each genotype, termed uphill ratio. Local sinks tend to have higher uphill ratios, while local peaks tend to have lower uphill ratios. We confirmed that genotypes with higher uphill ratios dominated at lower expression levels, while genotypes with lower uphill ratios dominated at higher expression levels (**Fig. 5F**). Importantly, even at the same expression levels, local peaks tended to have negative mutational changes, whereas local sinks tended to show positive changes. The fitness landscape was then modeled by adding the basal fitness, *F*_*b*_, to the expression levels multiplied by 1 − *F*_*b*_, where *F*_*b*_ ranged from 0 to 1 (**Fig. 5C, G**). That is, expression level is under positive directional selection in our model, where strength of selection pressure depends on *F*_*b*_. A smaller *F*_*b*_ corresponds to a steeper fitness landscape, while a larger *F*_*b*_ results in a flatter fitness landscape. This simple model assumes a positive linear relationship between expression level and fitness (**Fig. 5H**), which was relevant to the experimental evidence in most cases (**supplementary fig. S9B**). Based on the fact that knockdown of essential genes tends to be more deleterious for cell growth than nonessential genes (**supplementary fig. S9A**), we assigned lower and higher values of *F*_*b*_ for essential and nonessential promoters, respectively. Thus, in our model, essential and nonessential promoters shared the same expression landscape but experienced different strengths of selection pressures. Using the constructed fitness landscape with different *F*_*b*_ values (0.3, 0.5, and 0.99), we performed an evolutionary simulation based on the standard Wright-Fisher model(Otto and Day 2007) with asexual haploid population without recombination and with reversible mutation. Initial genotypes were randomly generated for each simulation. To mimic ongoing evolution of real organisms, the evolutionary simulation was performed for relatively short generations. In practice, the most abundant mutants within the evolved populations after 30 generations were isolated and subjected to measurement of mutational changes (**Fig. 5 G,H**). Despite the short generations, we confirmed that the evolved isolates in steeper fitness landscapes exhibited higher expression levels (**Fig. 5I**), which is consistent with the designed selection pressure. Comparing the mutational changes between different *F*_*b*_ values, we found that the mutants that evolved with smaller *F*_*b*_ showed negatively larger mutational changes, even at the same expression levels (**Fig. 5H**). These results suggest that higher activity and vulnerability in essential promoters are linked, and both can emerge simultaneously through stronger positive selection acting on higher expression levels of downstream essential genes.

**Fig. 5:**
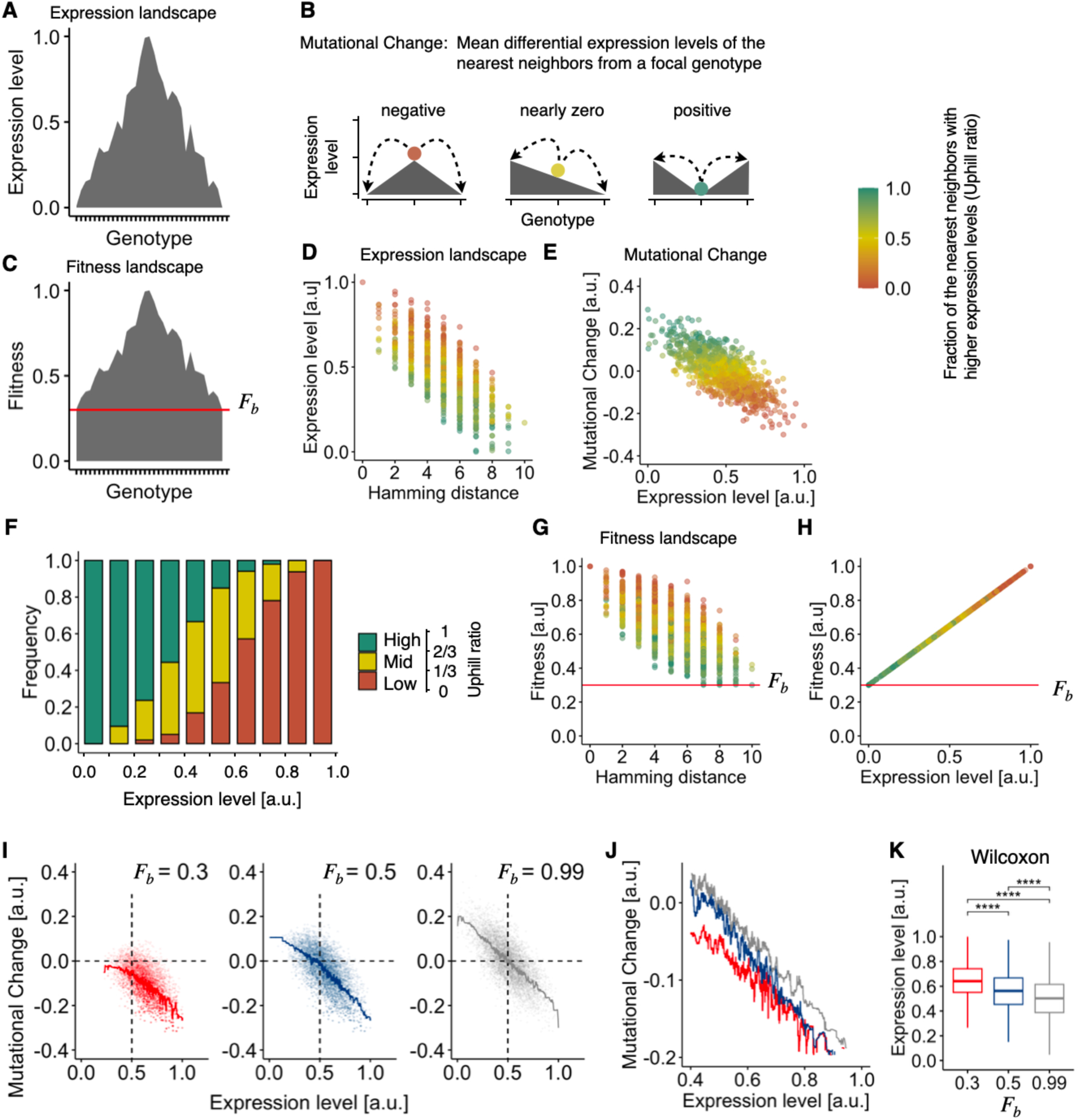
Numerical simulation of promoter evolution on a ragged fitness landscape. (**A**–**C**) Schematics of landscapes of expression levels (**A**, **B**) and fitness (**C**). (**A**) Expression landscape was constructed by the rough Mt. Fuji (RNF) model. (**B**) Mutational change of each genotype was calculated as the mean of differential expression levels of the nearest neighboring genotypes (arrow heads, mutant types) from the focal genotype (colored circles, wild type). The nearest neighbors differ from the focal genotype by one Hamming distance. The color bar represents the fraction of the genotypes with higher expression levels than the focal genotype among the nearest neighbors, termed uphill ratio. (**C**) Fitness landscape was modeled by adding basal fitness (*F*_*b*_, red line) to the expression landscape and normalizing it. (**D**, **E**) Representatives of expression landscape (**D**) and mutational change (**E**). Genotype was modeled as 10-bit binary numbers. Hamming distance represents the distance from the genotype with the global optima in expression level and fitness. (**F**) Relationship between expression level and uphill ratio. The uphill ratio was equally divided into three classes (Low, Mid, and High). Expression level was also equally divided into ten classes. Frequency of each class of uphill ratio was calculated as the number of corresponding genotypes divided by the total number of genotypes within the class. (**G**, **H**) Representatives of fitness landscape (**G**) and relationship between fitness and expression level (**H**) (*F*_*b*_= 0.3). (**I**) Mutational changes of the evolved isolates obtained through evolutionary simulation using the standard Wright-Fisher model. The basal fitness was varied by mimicking strong positive (*F*_*b*_ =0.3, left), moderate positive (*F*_*b*_ =0.5, middle), and nearly neutral selection (*F*_*b*_ =0.99, right) acting upon higher expression levels. Ten independent expression/fitness landscapes were generated for each condition of basal fitness. In each fitness landscape, 600 independent evolutionary simulations were examined where initial genotypes were randomly selected. Dots represents the most abundant mutants within the population after 30 generations in each run. Solid lines represent the running medians. Dashed lines are guides for the eye. (**J**) Enlarged figure of the running medians in (**I**). (**K**) Expression level of the evolved isolates. Asterisks represent significance levels of the adjusted p-value (the BH method) in the Wilcoxon test (****: p ≤ 0.0001).

Despite that our simple linear model basically agreed with the known relationship between expression level and growth fitness in most cases (**supplementary fig. S9**), an alternative model might be suitable for some genes such as *yidC* (**supplementary fig. S9B**). These genes exhibited almost no changes in growth fitness in response to small knockdown level, while growth fitness gradually decreases with increases in knockdown level beyond certain knockdown levels. To deal with such biphasic cases, we constructed the linear-plateau model (**supplementary fig. S9**). In short, the linear-plateau model has two phases: a positive linear phase and a flat plateau phase, where the joint point of the two phases is characterized by *E*_*p*_ ranging from 0 to 1 (**supplementary fig. S10D**). A larger *E*_*p*_ corresponds to a higher optimal expression level. Using the linear-plateau model with different *E*_*p*_ values, we performed an evolutionary simulation by following the same procedure as the linear model (**supplementary fig. S10E, F**). We found that the evolved isolates under the larger *E*_*p*_ exhibited higher expression levels (**supplementary fig. 10G**) and negatively larger mutational changes, even at the same expression levels (**supplementary fig. 10F**). These results seemed to be relevant by considering neutral evolution prevalent in the fitness landscapes with the smaller *E*_*p*_. These results highlighted an impact of optimal expression level on activity and vulnerability in promoters, suggesting that higher optimal expression levels could also contribute to higher activity and mutational vulnerability in essential promoters. Overall, our findings imply that both higher optimal expression and stronger directional selection could contribute to mutational vulnerability in expression levels for essential genes. Although the expression of essential promoters tended to decrease due to mutations (**Fig. 4E**), a few essential promoters with lower mean expression levels exhibited neutral to positive log_10_-fold changes in expression response to mutations. A prominent example was the essential promoter controlling *tilS*, which exhibited the lowest expression level among wild-type essential groups (**supplementary Table S2**). This promoter showed a positive value in log_10_-fold change in response to mutations, indicating the frequent occurrence of mutants with higher expression levels than wild-types. These essential promoters might suggest weaker directional selection or lower optimal expression levels for the downstream essential genes.

Consistent results were also obtained using the NK model(Kauffman and Weinberger 1989) for rugged expression landscapes (**supplementary figs. S11 and S12**, for the linear and linear-plateau models, respectively). The NK model, similar to the RMF model, frequently provided sinks (genotypes with higher uphill ratios) at lower expression levels and peaks (genotypes with lower uphill ratios) at higher expression levels (**supplementary fig. S11C**). However, compared to the RMF model (**Fig. 5F**), sinks were relatively rare at middle to high expression levels (beyond 0.3 in expression level). Interestingly, peaks were also rare at middle expression levels (0.3–0.6 in expression level) in the NK model. That is, genotypes with moderate uphill ratios were relatively frequent at middle to high expression levels. The low frequency of sinks and peaks indicates fewer genotypes with higher or lower mutational changes. Consequently, the difference in mutational changes between different genotypes at the same expression levels was relatively small in the NK model. Therefore, the difference in mutational changes between different *F*_*b*_ (or *E*_*p*_) values in the NK model was small relative to the RMF model (**supplementary figs. S11G**,**12F**). Despite these minor differences, both expression landscapes showed consistent outcomes for the linear (**Fig. 5J**,**K** and **supplementary fig. S11G**,**H** for the RMF and NK model, respectively) and linear-plateau models (**supplementary fig. S10F**,**G** and **supplementary fig. S12F**,**G** for the RMF and NK model, respectively). These results support the broad applicability of our key assumptions underlying rugged expression landscapes, suggesting the robustness of the outcomes derived from our simple models.

## Discussion

Transcriptional variability differs between genes. Using public transcriptome datasets for *E. coli*, we found that essential genes exhibited lower transcriptional variability across different mutants and environmental conditions than nonessential genes. To explore the underlying mechanism, we focused on the promoter regions. We quantified the changes in promoter activity changes in response to mutations by constructing mutant libraries of natural and synthetic promoters. As a result, essential promoters did not exhibit a significant difference in mutational variability compared with nonessential and synthetic promoters. Unexpectedly, we found that essential promoters showed remarkable vulnerability to mutations, where variants with higher activities than wild-type were rare and those with lower activity were abundant. These results imply that essential promoters have evolved to reach local optima in terms of promoter activity through past selections for higher expression levels of downstream essential genes. Our numerical evolutionary simulation confirmed the relevance of this scenario. Overall, our results suggest the evolutionary impact of essentiality on the directionality of the mutational variability of promoters and imply that the directional bias in mutational variability of promoter regions might constrain further evolution of expression levels of essential genes toward higher expression levels.

There are three well-known evolutionary scenarios for the emergence of mutational robustness in many biological systems(de Visser, et al. 2003), namely adaptive(Waddington 1942), congruent(Meiklejohn and Hartl 2002), and intrinsic(Siegal and Bergman 2002) scenarios. The adaptive scenarios predict that essential traits have evolved to reduce mutational variability and minimize the emergence of harmful mutants through direct selection for mutational robustness. The congruent scenarios predict that mutational robustness is a byproduct of evolution toward low variability against environmental perturbations. Contrary to these two scenarios assuming adaptive evolution toward low variability, the intrinsic scenarios predict that the mutational robustness of the traits emerges spontaneously and inherently along with the emergence of traits without selection against variability. In the case of promoter activity, an intrinsic scenario corresponds to the mutational robustness of promoters emerging spontaneously with the emergence of promoter activity without selection for mutational robustness, which corresponds to the mutational variability of synthetic promoters. Our results demonstrated that both adaptive and congruent scenarios failed to explain the observed mutational variability in natural promoters because essential promoters showed similar mutational variability to the other promoters, and the natural promoters exhibited no significant positive correlation with the transcriptional variability of downstream genes against different environmental perturbations. We note that the adaptive scenario might explain the higher expression levels, not transcriptional variability, of essential genes (**Fig. 1C, G**), which might minimize the risk of mutational knock down for essential genes to a lethal level. In contrast, the intrinsic scenario explained the observed mutational variability because the natural promoters showed similar mutational variability to that of synthetic promoters. Thus, the mutational variability of essential promoters is unlikely to be a consequence of adaptive evolution toward robustness against genetic and environmental perturbations.

Why did essential promoters show no less mutational variability than other promoters, despite the fact that the reduction in expression levels of essential genes is more likely to be harmful to cell growth than nonessential genes(Cui, et al. 2018)? One possible reason is that promoter sequences with significant mutational robustness are rare, although several possible mechanisms underlying mutational robustness in promoter regions are hypothesized(Payne and Wagner 2015). In other words, the expression landscape of the promoters might be rugged everywhere as we modeled in **Fig. 5A**. In fact, synthetic promoters derived from different random sequences achieved higher promoter activities equivalent to natural promoters, which was consistent with previous observations(Wolf, et al. 2015; Yona, et al. 2018). A second possible reason is that selection for higher expression levels on a rugged fitness landscape (**Fig. 5C**) prevents the essential promoters at local optima from traversing the expression landscape to reach genotypes with higher mutational robustness. Consistently, essential promoters showed biased mutational effects, where most mutations in essential promoters tended to reduce promoter activity (**Fig. 4C**). Interestingly, this scenario agreed with the predictions from a machine-learning model based on an empirical expression landscape of yeast promoters(Vaishnav, et al. 2022). Thus, our results suggest several plausible hypotheses that may be shared by different organisms. Ideally, future studies should explore evolutionary scenarios for diverse promoters that were not tested in this study, and validate the hypothetical landscape of promoters more thoroughly. The increase of the sample set of promoters would also compensate for our small sample sets and would enhance power to detect a potential subtle difference in mutational variability between different promoter groups.

In this study, we focused only on mutations in promoter regions to explore the mechanism underlying the lower transcriptional variability in essential genes. As a result, we found that mutants with higher promoter activity were relatively rare for essential promoters. However, this result does not exclude the contribution of any other mechanism underlying the observed transcriptional variability in response to genome-wide perturbations (**Fig. 1)**. Notably, *trans*-acting mechanisms, such as transcriptional regulators, may also play a role. As it is, the previous studies reported that genes exhibiting greater transcriptional variability across diverse environmental and genetic perturbations tend to be regulated by a larger number of unique transcriptional regulators in *E. coli*(Tsuru and Furusawa 2024) and yeast(Landry, et al. 2007). A higher number of regulators provides more *trans*-mutational target sites for the regulated genes, thereby enhancing transcriptional variability in response to genomic perturbations. Based on this evidence, it is plausible that essential genes have less transcriptional regulation. To support this hypothesis, we used data from RegulonDB and EcoCyc to compare the number of regulations between essential and nonessential genes (**supplementary fig. S13**). First, we found a smaller fraction of essential genes (90 of 248 genes, 36%) lacking known transcriptional regulators than nonessential genes (1976 of 3995, 49%) (**supplementary fig. S13A**). This finding aligns with the positive activation bias values observed (**Fig. 4F**), probably attributable to the rich activation by sigma factors aimed at enhancing expression levels. On the other hand, among the genes regulated by at least one transcriptional regulator, we observed that essential genes received fewer transcriptional regulations than nonessential genes (**supplementary fig. S13B**), supporting this hypothesis. This observation is consistent with previous findings in yeast(Bilu and Barkai 2005). Thus, our study underscores the dual contributions of both *cis*- and *trans*-acting mechanisms including their epistatic interactions in constraining the transcriptional variability of essential genes.

## Materials and Methods

### Bacterial strains and medium

We used *E. coli* DH5α Electro-Cells (Takara Bio, Japan, 9027) to construct the pUA66- mCherry plasmid and its derivatives. LB medium (LB Broth, Miller, BD Difco, USA, 244620) supplemented with 25 µg/mL kanamycin (Kanamycin Sulfate, Wako, Japan, 115-00342), referred to as LB/Km medium, served as the selective medium. The SOC medium, derived from SOB medium (SOB Medium, BD Difco, USA, 244310) supplemented with 22 mM Glucose (Wako, Japan, 049-31165), was used for recovery culture pos-transformation. The *E. coli* K-12 MG1655 strain served as the host for the constructed plasmids and was employed for measuring promoter activity via flow cytometry. Bacterial cells for flow cytometry were cultured in a synthetic medium, a derivative of M9 minimal medium (pH 7.2), supplemented with 47.7 mM Na_2_HPO_4_ (Disodium Hydrogenphosphate 12-Water, Wako, Japan, 196-02835), 22.0 mM KH_2_PO_4_ (Wako, Japan, 164-22635), 8.55 mM NaCl (Wako, Japan, 191-01665), 9.35 mM NH_4_Cl (Wako, Japan, 014-03005), 11 mM Glucose, 1 mM MgSO_4_ (Magnesium Sulfate Heptahydrate, Wako, Japan, 138-00415), 50 µM CaCl_2_ (Wako, Japan, 038-24985), and 3.8 µM thiamin (Thiamin Hydrochloride, Wako, Japan, 201-00852), with supplementation of 25 µg/mL Kanamycin when required.

### Selection of natural promoters

The *E. coli* Promoter Collection(Zaslaver, et al. 2006) (Horizon Discovery Ltd., UK, PEC3877) was used to obtain the promoter regions of the natural promoters in *E. coli*. Initially, we filtered out the promoters with low activities that were near our detection limit based on the promoter activities of this library measured by Silander et al(Silander, et al. 2012). This filtration excluded 12% and 17% of essential and nonessential promoters, respectively. Subsequently, 115 promoters were randomly selected from this subset library carrying the pUA66 plasmid. The DNA sequences of these promoter regions were verified using Sanger sequencing (Azenta Life Sciences, USA) and compared with the reference genome sequence of *E. coli* K-12 MG1655(Blattner, et al. 1997) (GenBank ID: U00096). Subsequently, 63 of the 115 strains without mutations in their promoter sequences were identified, and plasmids were extracted from these colonies. Among these, 56 strains were randomly selected for the subsequent experiments (**supplementary table S2**). Each promoter was denoted by the “NP” (natural promoter), followed by the name of the ORF present in the immediate downstream sequence in the genome. For example, NPmetG represents the promoter regions upstream of the *metG* gene on the chromosome.

### Plasmid construction

Plasmid pUA66 was modified by replacing the promoter regions of *gfp*, flanked by the BamHI and XhoI restriction sites, with a red fluorescence protein (mCherry) expression cassette. This cassette comprised dual terminators RNAI and TSAL(Reynolds, et al. 1992), promoter PLtetO- 1(Lutz 1997), Shine-Dalgarno sequence SD8(Ringquist, et al. 1992) and codon-optimized mCherry(Dunlop, et al. 2008) coding sequences. The terminators were obtained from pNS2-VL(Dunlop, et al. 2008)(Addgene, Plasmid #26756) and the assembled sequences of the other elements were commercially synthesized (Fasmac Co., Ltd., Japan). The assembled cassette was inserted between the BamHI and XhoI recognition sequences of the pUA66 plasmid in the direction opposite to that of *gfp* (**supplementary fig. S2**), yielding plasmid pUA66-mCherry. PCR amplification was performed using KOD One PCR Master Mix Blue (Toyobo, Japan, KMM-201) and pUA66-mCherry as a template with primers A (GCTTTCCAGGGATCCTCTAGATTTAAGAAGG) and B (GAGCAGTACTCTCGAGGTGAAGACGAAAGG) to include all sequences except for the mCherry expression cassette. The PCR product was column purified using the FastGene Gel/PCR Extraction Kit (NIPPON Genetics, Japan, FG-91302). The purified PCR product was double-digested with BamHI (Takara Bio, Japan, 1010A) and XhoI (Takara Bio, Japan, 1094A) overnight at 37 °C, followed by column purification. The digested product is referred to as the vector.

### Construction of promoter library comprising of random sequences

The construction of the synthetic promoters was based on a previous study(Wolf, et al. 2015). Single-stranded oligonucleotides (CCTTTCGTCTTCACCTCGAG-(N200)- GGATCCTCTAGATTTAAGAAGG) were synthesized by Eurofins Genomics (Ebersberg, Germany). The 5’ and 3’ ends of the oligos contained XhoI and BamHI restriction sites, respectively, and were homologous to primers C (CCTTTCGTCTTCACCTCGA) and D (CCTTCTTAAATCTAGAGGATCC), respectively (**supplementary fig. S2**). A 200 bases region of random nucleotides was present between them. PCR amplification using primers C and D was performed using a mixture of single-stranded oligos as a template to create double-stranded DNA. The PCR products were column purified and double digested using BamHI and XhoI for 3 h at 37 °C, followed by column purification. Thus, an insert for the following ligation reaction was obtained. The ligation reaction was performed using T4 DNA Ligase (Takara Bio, Japan, 2011A) overnight at 16 °C involved mixing the vector and insert at a molar ratio of 1: 3. The resulting ligation product solution was column purified and DNA was extracted using 20 µL of sterile water. All purified ligation product solutions were added to 50 µL of DH5α Electro-Cells and electroporated at 1800 V (Eporator, Eppendroph, Germany, 4309000027). Transformed cells were incubated at 37 °C for 1 h in 1 mL of SOC medium, followed by overnight incubation at 37 °C in 1 mL of LB medium supplemented with 2 µL of 25 mg/mL kanamycin solution.

### Selection of synthetic promoters using flow cytometry

Green and red fluorescence, as well as forward scatter of the cells in the DH5α-transformed random promoter library were measured using a flow cytometer (FACSAria III, BD, USA). Gates were set for the green and red channels to exclude cells with low GFP and high mCherry fluorescence, respectively. Subsequently, 10,000 cells without red fluorescence but with green fluorescence within the selected range (**Fig. 2B**) were collected by cell sorting, according to the manufacturer’s instructions. The sorted cells were spread on LB/Km agar plates and incubated overnight at 37 °C. Bacterial colonies were randomly picked and their inserted synthetic promoter regions were sequenced by Sanger sequencing. Consequently, 16 synthetic promoters without improper mutations at the restriction sites were identified and named SP1–16, (**supplementary table S1**).

### Random mutagenesis using error-prone PCR

Error-prone PCR was conducted using 0.1 ng of each promoter region as a template, with 35 cycles of amplification performed using the GeneMorph II Random Mutagenesis Kit (Agilent Technologies, USA, 200550) and primers C and D. The resulting PCR products underwent gel purification and were double digested with BamHI and XhoI for 3 hours at 37 °C, followed by column purification to create mutant inserts. Ligation was performed overnight at 16 °C using T4 DNA Ligase, with the vector and mutant inserts mixed at a molar ratio of 1:3. The ligation reaction solution was column purified, and DNA was extracted using 20 µL of sterile water.

### Construction of mutant promoter library

The purified ligation product solution was added to 50 µL of DH5α Electro-Cells and electroporated at 1800 V. Transformed cells were cultured at 37 °C for 1 h in 1 mL of SOC medium and a portion of the culture was plated on LB/Km agar medium for overnight growth at 37 °C. Bacterial colonies were counted to determine the transformation efficiency. If the transformation efficiency exceeded 10^5^ cells/reaction, the next step was performed; otherwise, the process was repeated. The transformed culture was diluted with 5 mL of LB/Km medium and grown overnight. Plasmid extraction from an overnight culture was performed using the FastGene Plasmid Mini Kit (NIPPON Genetics, Japan, FG-90502). The obtained plasmid solution was added to 50 µL of MG1655 electrocompetent cells and electroporated at 1800 V. Transformed cells were cultured at 37 °C for 1 h in 1 mL of SOC medium, and a portion of the solution was plated on LB/Km agar medium for overnight growth at 37 °C to count the number of colonies and determine transformation efficiency. If the transformation efficiency exceeded 10^5^ cells/reaction, the next step was performed; otherwise, the process was repeated. A sterile 2 mL 96 well deep well plate (Greiner Bio-One, Austria, 780271) was used to add 1 mL of the synthetic medium to each well, followed by the addition of 10 µL of the overnight culture containing the MG1655 transformants. The bacterial cells were grown at 37 °C with shaking at 800 rpm until confluent growth was achieved. Subsequently, another plate was prepared with 1 mL of fresh synthetic medium added to each well, and 10 µL of the confluently grown cultures was diluted into each well. The cultures were grown for 2.5 h at 37 °C with shaking at 800 rpm and were used as mutant libraries for flow cytometry.

### Verification of PCR random mutagenesis

The number of mutations introduced into the promoters by error-prone PCR was determined using Sanger sequencing. Three to four colonies were randomly selected from six randomly selected mutant libraries, including essential and nonessential promoters (**supplementary table S2**). Colony PCR was performed using KOD One PCR Master Mix Blue with primers E (TTACTTTGCAGGGCTTCCCAA) and F (CCAGTCTTTCGACTGAGCCT) to amplify the regions including the promoter regions (**supplementary fig. S2**). PCR products were column purified and verified by Sanger sequencing. The average mutation rate was calculated to be 6.2 mutations/kb. Mutation rates and mutational spectrum for each promoter group were shown in **supplementary fig. S8**.

### Construction of wild-type promoter control group

To ensure accurate measurement of GFP fluorescence from cells with wild-type promoters, wild-type plasmids were transformed into MG1655 cells by electroporation. Transformed cultures were plated on LB/Km agar medium and incubated overnight, and the resulting colonies were cultured in LB/Km medium. Cell cultures of these control groups in the synthetic medium were prepared and used as wild types for flow cytometry.

### Flow cytometry

Bacterial cells were dispensed in M9 buffer prior to analysis using a flow cytometer (BD, USA, FACSAria III). Fluorescence intensity from GFP (GFP FI) and mCherry (mCherry FI) was measured using a 488 nm laser with 515–545-nm emission filter, and a 561 nm laser with 563– 588-nm emission filter, respectively. The following PMT voltage settings were used: forward scatter (FSC), 300; side scatter (SSC), 340; green fluorescence, 700; red fluorescence, 600. The events exceeding the defined thresholds of FSC (200) and SSC (200) were recorded. Tota1,000,000 events were measured per library. The MG1655 strain was used as a control for autofluorescence of GFP FI.

### Data analysis of flow cytometry

FCS files were converted to CSV format using the flowCore(Ellis, et al. 2024) package in R(R Core Team 2023). Subsequent analyses were conducted using custom R scripts. To compare green fluorescence among cells in similar physiological states, a narrow gate for the forward scatter channel was set to capture 40% of the events, including the most frequent values for MG1655 (**supplementary fig. S5A**). The top 0.5% value of the green fluorescence intensity distribution of MG1655 within the gate was determined as the autofluorescence intensity (**supplementary fig. S5B**). For the events of wild types and mutant libraries, the autofluorescence intensity was subtracted from the green fluorescence intensity of the events within the gate. Events with a negative green fluorescence intensity after subtracting the autofluorescence intensity were excluded from the analysis. In addition, events with a red fluorescence intensity greater than 10^3^ [a.u.] were regarded as the cells carrying pUA66- mCherry and were excluded from the analysis. Clonal cell populations exhibit huge cell-to-cell heterogeneity in gene expression even under identical environmental conditions, resulting in long-tailed or log-normal shaped distributions in protein numbers(Tsuru, et al. 2009; Taniguchi, et al. 2010). This stochastic nature often results in rare cells exhibiting very highly expression levels within cell populations, which greatly influences variance or standard deviation of protein number distributions. To ensure robust measurement of variation in fluorescent distributions, the green fluorescence intensity after subtracting the autofluorescence intensity was log_10_-transformed and used to calculate the means and standard deviations. We note that this log-transformation might limit detection of possible small differences in mutational variability in promoter activities between different promoter groups.

### Classification of promoter based on essentiality of transcribed unit

Gene essentiality was based on a previous study(Goodall, et al. 2018), which identified 248 genes designated as essential across different datasets. In bacteria, clusters of genes are controlled by the same promoter sequences and are transcribed as single mRNAs, termed transcription units. Promoters controlling at least one essential gene in the transcription units were defined as essential promoters (**Fig. 2A**). The known transcription units were retrieved from the RegulonDB(Tierrafria, et al. 2022) database. Of the 56 promoters, 18 were classified as essential and 38 as nonessential as listed in **supplementary table S1**.

### Estimation of transcriptional variability

Transcriptional variability of each gene in *E. coli* in response to different genetic perturbations was estimated using known transcriptome profiles obtained from six independent mutants derive from MG1655 cultured under a single environmental condition, termed the Mut dataset(Tsuru and Furusawa 2024). These mutants were subjected to mutation accumulation (MA) experiments to accumulate genomic mutations via genetic drift(Halligan and Keightley 2009; Tsuru, et al. 2015). In addition, to quantify the transcriptional variability in response to different environmental conditions, known transcriptome profiles of MG1655 cultured under 76 different environmental conditions(Sastry, et al. 2019; Rychel, et al. 2021; Lamoureux, et al. 2022), termed the Env dataset(Tsuru and Furusawa 2024), were utilized. The expression levels were determined by log_2_-transformation and quantile normalization. The means and standard deviations in expression levels for each gene across different genetic perturbations and environmental conditions were quantified. To compensate for the mean-dependency of the standard deviations (**Fig. 1B**,**G**), the DM values for each gene were calculated as the vertical distance of each standard deviation from a smoothed running median of the standard deviations for each dataset, as detailed previously(Tsuru and Furusawa 2024). The resulting DM values were termed DM_mut_ and DM_env_ for the Mut and Env datasets, respectively. The DM values were used as measures of transcriptional variability in response to genetic and environmental perturbations.

### Phylogenetic analysis of promoter sequence of length 73 bases

Synthetic promoters underwent a blastn(Altschul, et al. 1990) search to confirm that there was no considerable similarity to any natural sequences (**supplementary note**). Posterior sequences of 73 bases were aligned using the msa(Bodenhofer, et al. 2015) package in R (Clustal Omega), where 73 bases represented the shortest length of a synthetic promoter. A phylogenetic tree was constructed using the Neighbor-Joining method.

### Phylogenetic analysis of core elements of sigma70 promoters

We identified sigma70 promoters from our natural promoters using RegulonDB. The core elements comprising the –10/–35 elements of these promoters were concatenated and subjected to phylogenetic analysis. The Levenshtein distance(Levenshtein 1966) between the focal elements and the consensus elements, which consist of TTGACA and TATAAT, was calculated. A phylogenetic tree was constructed as previously described.

### Identification of the number of transcriptional regulators regulating the natural promoters

Reported regulatory interactions between genes were compiled using all interactions with experimental evidence from RegulonDB and EcoCyc(Keseler, et al. 2021) for both transcription factors and sigma factors. The mode of regulation was classified as either activation or inhibition, and the mode of regulation by sigma factors was classified as activation.

### Numerical simulation of promoter evolution

The promoter genotype, *g*, was modeled by a bit string comprising either zeros or ones of length N=10 (e.g., 1110001100). The expression level, *E*, was defined by the Rough Mt. Fuji (RMF) model(Aita, et al. 2000; Neidhart, et al. 2014) as:

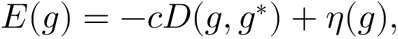

where *c* is a scaling coefficient equal to one in this study, *g*^∗^ is the genotype with the highest expression level before adding noise, *D*(*g*, *g*^∗^) is the Hamming distance between *g* and *g*^∗^, and *η* is an independent and identically distributed random variable. A standard normal distribution with a mean equal to zero and a standard deviation equal to one for *η* was used to generate a rugged expression landscape. To construct an expression landscape, *g*^∗^ was first chosen at random from the genotype space. *E*(*g*) was subsequently calculated using the above equation. The expression landscape was then normalized using the minimal and maximal values of *E*(*g*) to range from zero to one, as follows:

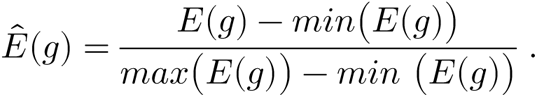

The fitness of a given genotype *g* was defined as:

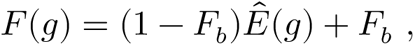

where 1≥*F*_*b*_≥0. *F*_*b*_ represents the basal fitness when the expression level is zero and determines the strength of the selection pressure acting on the expression level. Different values of *F*_*b*_ (0.3, 0.5, and 0.99) were used to demonstrate the evolution under strong positive selection, moderate positive selection, and nearly neutral evolution, respectively. It was assumed that essential genes underwent strong positive selection on their expression levels, while nonessential genes underwent relatively moderate selection for their expression levels, based on the empirical fact that knock-down of essential genes was more harmful for cell growth than nonessential genes(Li, et al. 2016). Ten independent fitness landscapes were constructed for each *F*_*b*_.

Evolutionary simulation on the fitness landscapes were performed using the standard Wright-Fisher model(Otto and Day 2007) assuming a haploid asexual population with a population size of 10^5^ and a mutation rate of 10^−3^ mutations per genotype per generation. The initial genotypes were randomly selected for each simulation, and 600 independent simulations were examined for each fitness landscape. For each generation, mutants with genotypes differing by one Hamming distance from the current population were generated at the mutation rate. The frequency of each genotype within the population was updated based on the fitness landscape. Subsequently, a new population for the next generation was formed by sampling mutants, with replacement, from the current population using a binomial distribution with the given population size. The most frequent mutants in a population after 30 generations were isolated in each simulation, resulting in 6,000 mutants for each *F*_*b*_. Mutational changes in expression levels were calculated for each isolate by averaging the expression levels of all nearest neighboring genotypes that differed by one Hamming distance from the focal isolate. Custom R scripts were used for the numerical simulations, modified from scripts constructed in a previous study(Obolski, et al. 2018; Song and Zhang 2021).

### Data visualization

All figures were generated in R and the following packages were used. Phylogenetic trees of the promoters were visualized using the ape(Paradis and Schliep 2019) and ggtree(Yu, et al. 2017) packages. Other illustrative plots were generated using the ggplot2(Wickham 2016) and ggpubr(Kassambara 2023) packages.

## Data Availability

Custom R scripts used in the manuscript are available at GitHub (https://github.com/tsuruubi/Variability_PromoterAct_E_coli).

The raw fcs data are available at Zenodo (wild types for https://doi.org/10.5281/zenodo.11108555; mutant libraries for https://doi.org/10.5281/zenodo.11108786).

## Acknowledgments

The authors thank Hiroyo Koike for technical assistance of Sanger sequencing. The authors also appreciate Dr. Olin Silander and Dr. Luise Wolf for fruitful technical advice on the construction of a synthetic promoter library. The pNS2-σVL plasmid was a gift from Dr. Michael Elowitz (Addgene, plasmid # 26756).

