## supplementary fig for "Promoters constrain evolution of expression levels of essential genes in *Escherichia coli*"

**I. Supplementary Figures**

**II. Supplementary Note**

**III. Supplementary References**

### I. Supplementary Figures

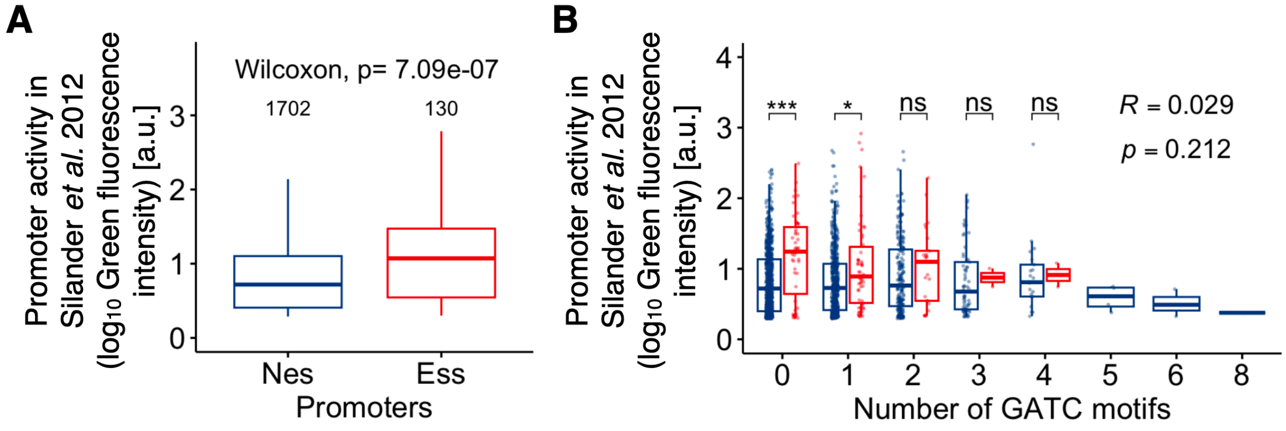


**Fig. S1: Promoter activities based on a public dataset**

(**A**) Promoter activities of *E. coli* natural promoters, originally reported by Silander et al(Silander, et al. 2012), were compared between nonessential and essential promoters. Promoters controlling at least one essential gene in the transcription units were defined as essential promoters, while those controlling only nonessential genes were defined as nonessential promoters. Promoter activity was measured by flow cytometry, with GFP transcriptionally fused to the plasmid copies of promoter regions. (**B**) Relationship between promoter activity and the number of GATC motifs in promoter regions. The lower and upper edges of the boxes in panels **A** and **B** represent the first (q1) and third (q3) quartiles, respectively. The horizontal lines in the boxes represent the medians (m). The whiskers from the boxes extend to the most extreme observed values inside inner fences, m±1.5(q3–q1). Pairwise Wilcoxon test was examined with a p-value adjustment (the BH method, ns: $p > 0.05$; *: $p \leq 0.05$; ***: $p \leq 0.001$). Spearman's R and p-values are shown in pane **B**.

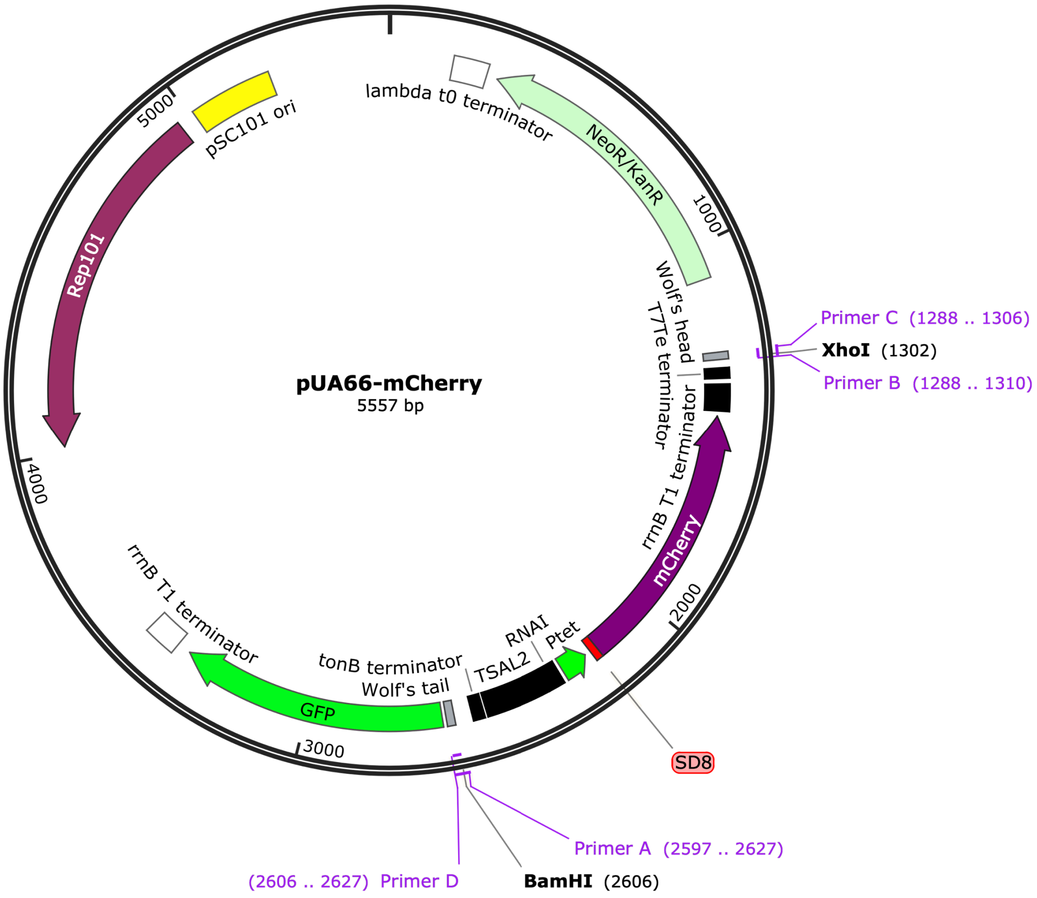


**Fig. S2: Map of the pUA66-mCherry plasmid**

Primers A–D are shown in purple. Grey blocks, labeled Wolf's head and tail, represent the homologous sites of the primers. The shorter region in-between Wolf's head and tail was replaced with the promoter regions of interest. This replacement was confirmed by the lack of red fluorescence derived from mCherry. Black blocks represent terminators.

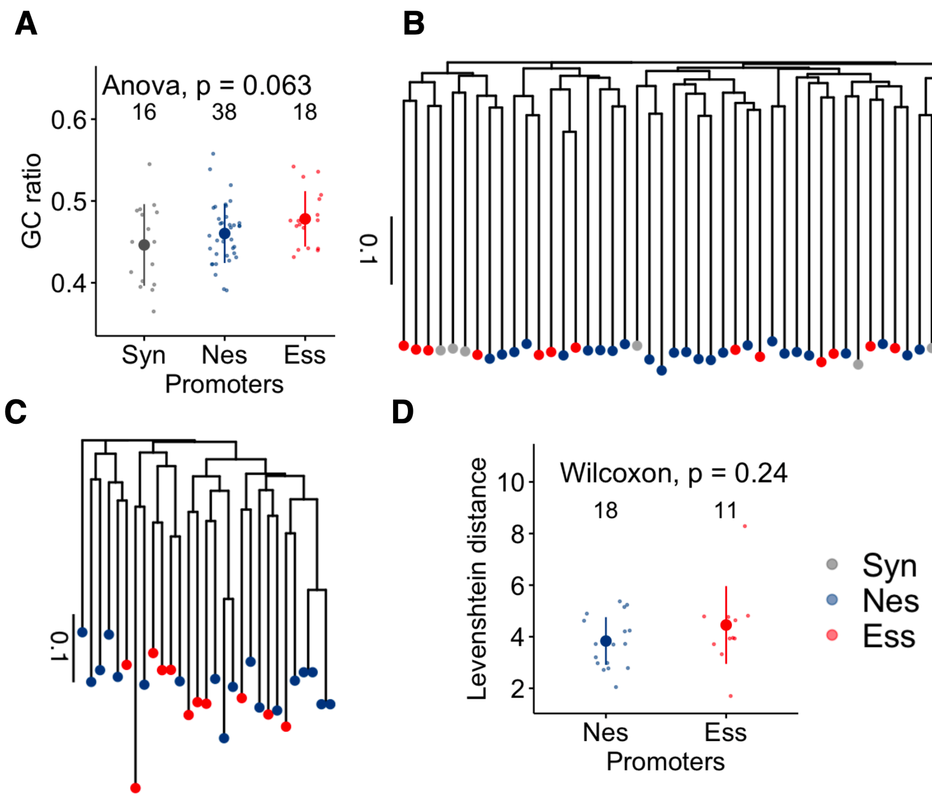


**Fig. S3: Sequence homology between synthetic and natural promotes.**

(**A**) GC content of three promoter groups (Syn, Nes, and Ess). (**B**) Phylogenetic tree of promoter regions using multiple sequence alignment (MSA) based on 73 bases from the posterior ends of the promoter regions. (**C**) Phylogenetic tree of the combined sequences of the –10 and –35 elements of the sigma70 promoter using MSA. The –35 and –10 elements of the promoters were identified from RegulonDB(Tierrafria, et al. 2022) and combined. (**D**) Edit distances of the –10 and –35 elements of sigma70 promoters are shown. The Levenshtein distance(Levenshtein 1966) between the combined sequences and concatenated consensus sequence (TTGACATATAAT) was calculated. The large dots, error bars, and inset numbers in panels **A** and **D** represent the mean, standard deviation, and number of genes, respectively.

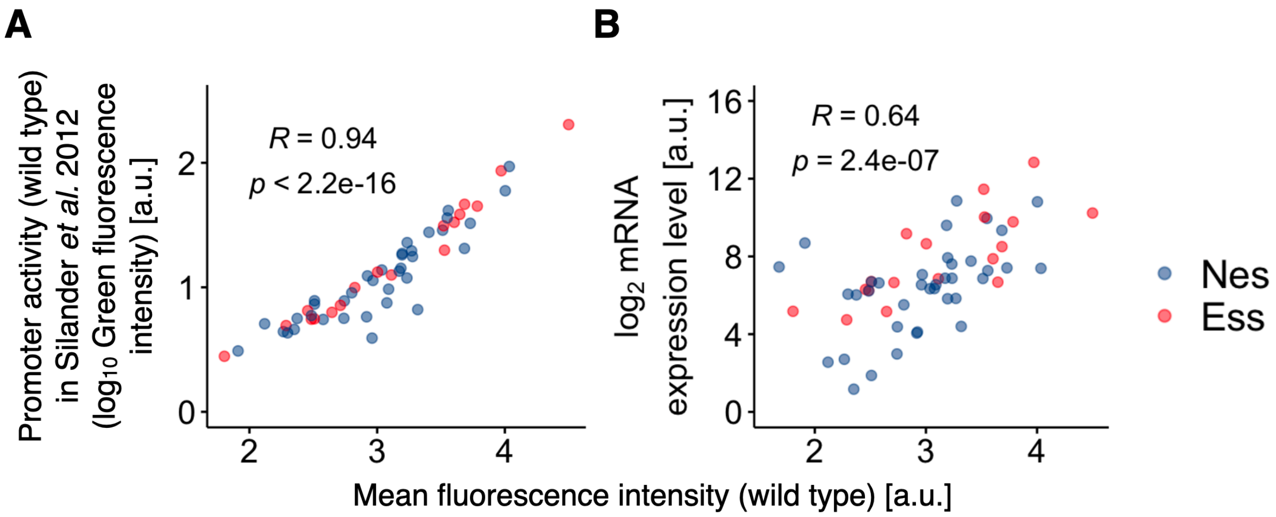


**Fig. S4: Consistency of measured promoter activities and their relationship with mRNA expression levels.**

(**A**) Correlation of wild-type promoter activity between two studies. The x-axis represents the promoter activities of the natural promoters measured in this study, while the y-axis represents the activities of the same promoters measured by Silander *et al*(Silander, et al. 2012). (**B**) Relationship between the promoter activities measured in this study and the mRNA expression levels of the genes downstream of the chromosomal copies of the promoters. The mRNA expression levels were obtained from the Env dataset (**Fig. 1E**), which represents the mean expression levels of the genes across different environmental conditions. Spearman's R and p-values are indicated.

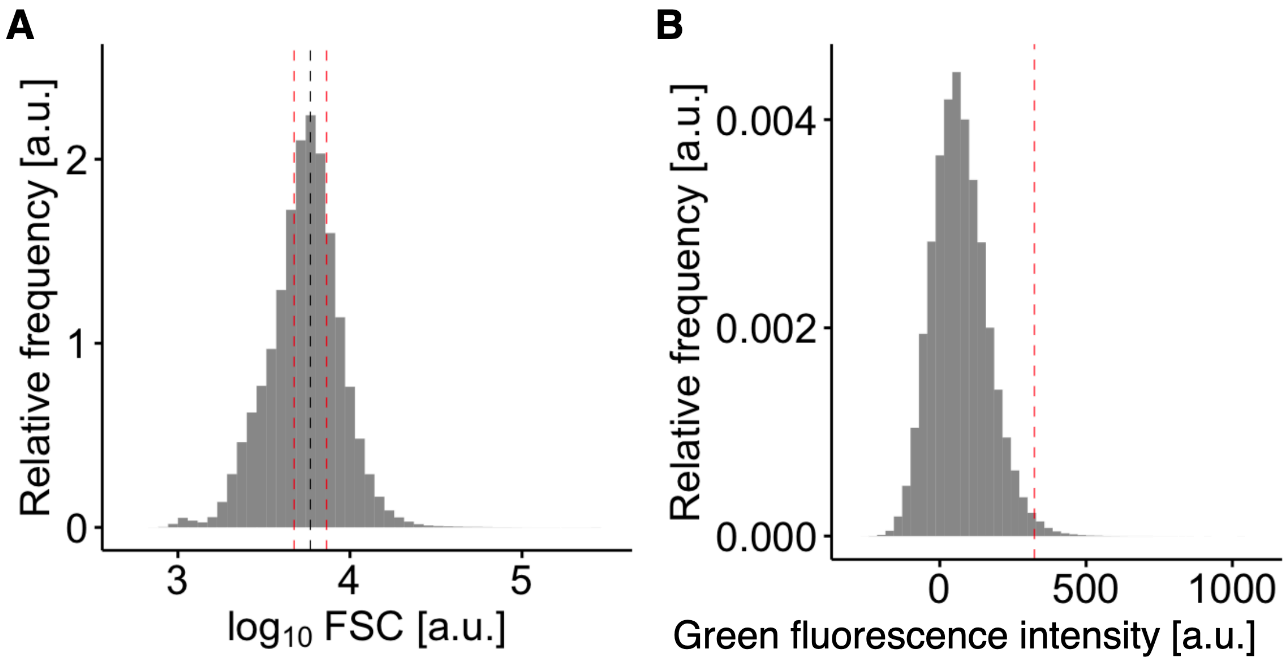


**Fig. S5: Forward scattering distribution and autofluorescence of the control stain.**

Forward scattering (FSC) distribution (**A**) and autofluorescence distribution (**B**) of MG1655. The black and red dashed lines in (**A**) represent the mode and gate for reducing the cell-size dependency of the green fluorescence, respectively. The red lines are defined to contain 20% of the total cells between the red and black lines. Consequently, 40% of the cells were subjected to the subsequent analyses. The red dashed line in (**B**) represents the top 0.5% of the autofluorescence distribution.

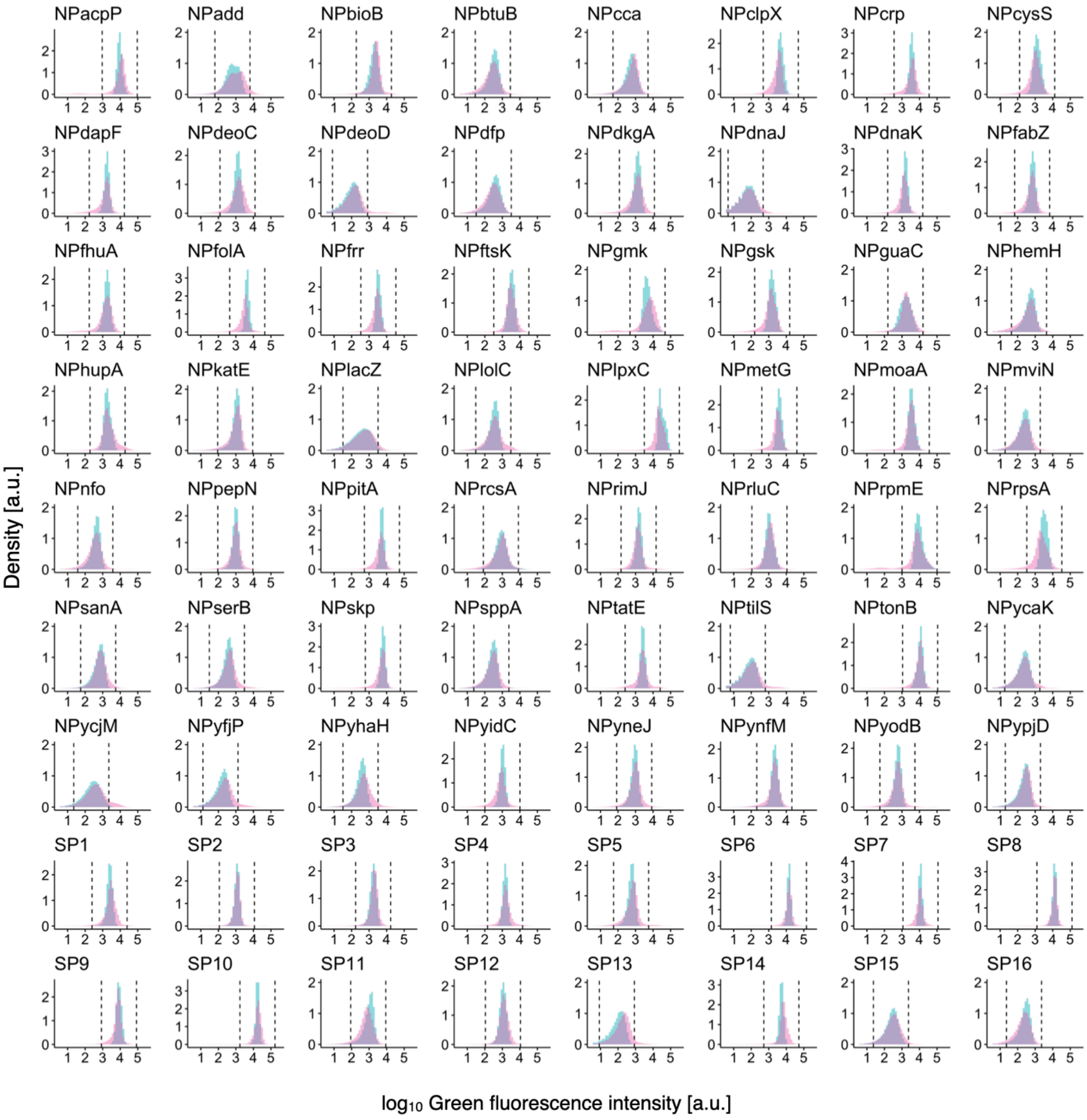


**Fig. S6 Green fluorescence distribution of all promoter libraries.**

Promoter names are indicated at the top of each panel. The cyan and pink distributions correspond to wild types and mutant libraries, respectively. The vertical dashed lines in each pane represent the lower (left) and upper (right) thresholds used for analysis in **supplementary fig. S7**.

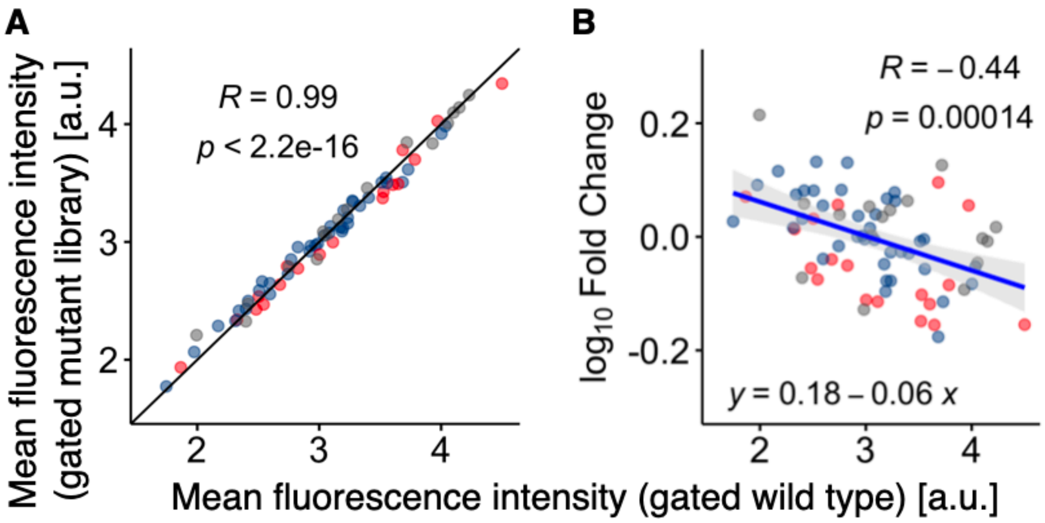


**Fig. S7 Mean expression changes of cells within the limited detection ranges.**

In principle, the negative correlation shown in **Fig. 4A** could be a technical biproduct of the asymmetry in detecting mutational changes in expression level, where a decrease in expression becomes more difficult to detect as the expression level decreases. To exclude this possibility, we calculated the mean expression levels for the cells within defined narrow gates shown in **supplementary fig. S6** (dashed lines). First, we calculated the mean expression levels for wild-type cells without gating. The lower and upper thresholds of the gates were subsequently defined by adding 1 a.u. to or subtracting 1 a.u. from the ungated means, respectively. Importantly, the gate size was fixed as 2 a.u. over expression levels. The mean expression levels were then calculated for the cells within the gates for the wild types and mutant libraries (**A**). (**B**) Relationship between the $\log_{10}$-fold change of mean expression levels, from gated wild types to gated mutant libraries, and the expression levels of gated wild types. The blue line represents a regression line. Spearman's R and p-values are shown in panels **A** and **B**. Grey, blue, and red dots in each panel represent synthetic (Syn), nonessential (Nes), and essential (Ess) promoters (defined in **Fig. 2**), respectively. The negative correlation in panel **B** supports that the negative correlation in **Fig.4B** was not due to the technical biproduct of our limitation of detecting a decrease in expression levels.

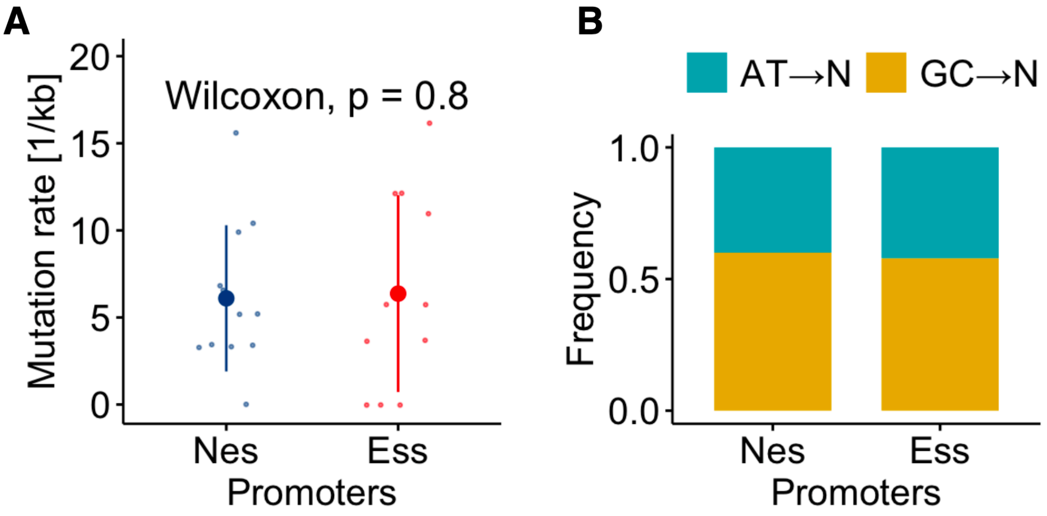


**Fig. S8: Mutation rates and spectrum in error-prone PCR.**

(**A**) Base-pair substitution rates of two promoter groups (Nes and Ess). Six randomly selected promoter libraries, three from essential promoters and the rest from nonessential promoters, were sequenced to detect mutations in the promoter regions. Three to four colonies were analyzed for each library (**supplementary Table S2**). The mutation rates were calculated from the number of substitutions divided by the length of promoter regions. (**B**) Mutational spectrum based on the identified base-pair substitutions (**supplementary Table S2**). The base-pair substitutions from AT to other base pairs (TA, GC, CG) and those from GC to other base pairs (CG, AT, TA) are colored in blue-green and orange, respectively. The frequency of each substation group (AT→N or GC→N) was calculated from the number of focal substitutions divided by the total number of substitutions within each promoter group (Nes or Ess).

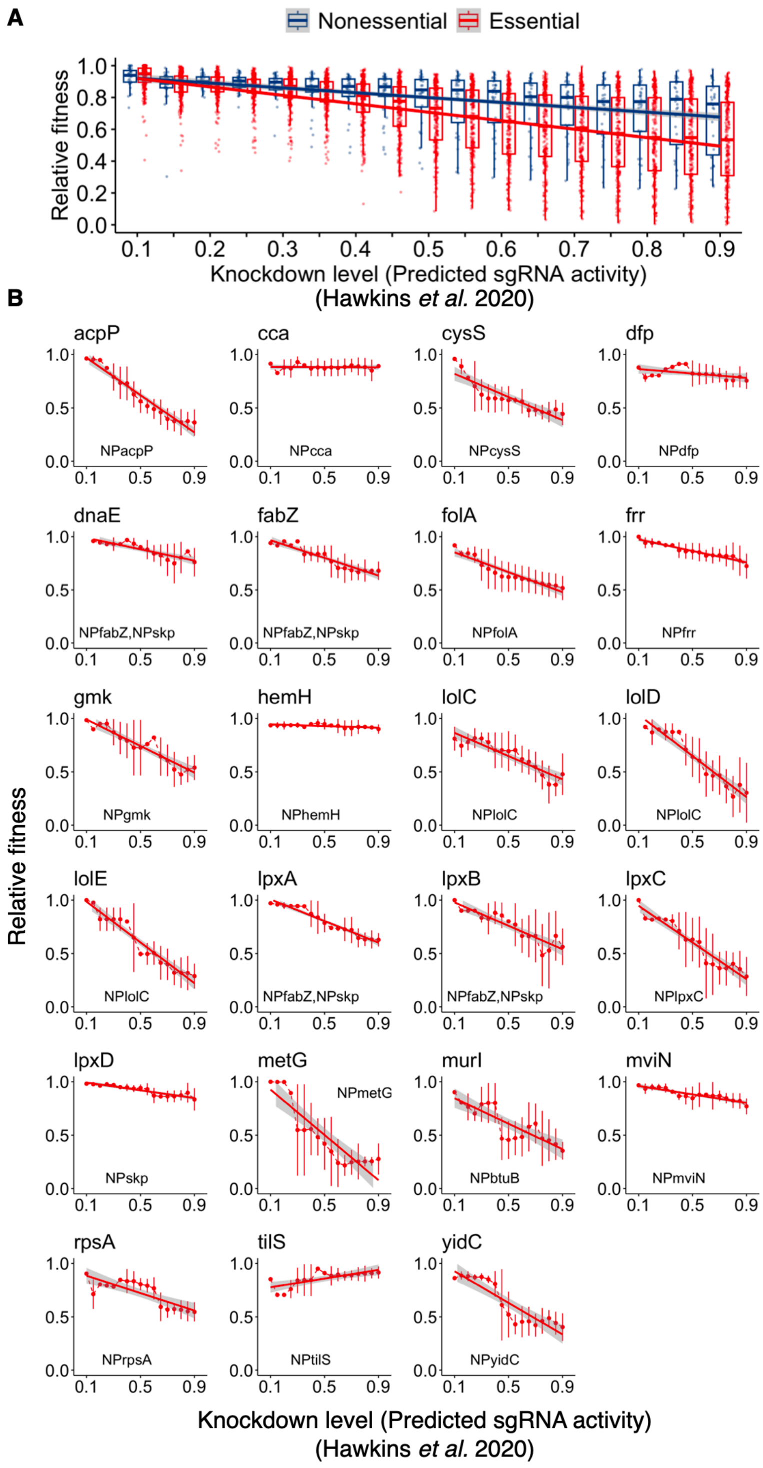


**Fig. S9: Relationship between growth fitness and knockdown of expression.**

(**A**) Relationship between growth fitness and knockdown level for 236 essential (red) and 34 nonessential (blue) genes in *E. coli*. The dataset was downloaded from Hawikins *et al*(Hawkins, et al. 2020). In brief, Hawikins *et al* performed knockdown using a CRISPRi system with mismatched sgRNAs for most essential genes. Knockdown level represents sgRNA activity predicted from a linear model based on a training dataset(Hawkins, et al. 2020). Higher knockdown levels represent lower expression levels. Relative fitness was based on a competitive growth experiment between non-targeting control library and essential-gene targeting library, representing the median fitness of the mismatched sgRNAs within each bin of sgRNA activity. Solid lines represent linear regression lines (red and blue for essential and nonessential genes, respectively). For consistency, gene essentiality was reassigned based on the definition used in the current study. (**B**) Relationship between growth fitness and knockdown level for 23 essential genes controlled by essential promoters used in the current study. Top and inset labels in each subpanel represent names of essential genes and essential promoters, respectively. Error bars represent the median absolute deviations(Hawkins, et al. 2020). Solid lines represent linear regression lines.

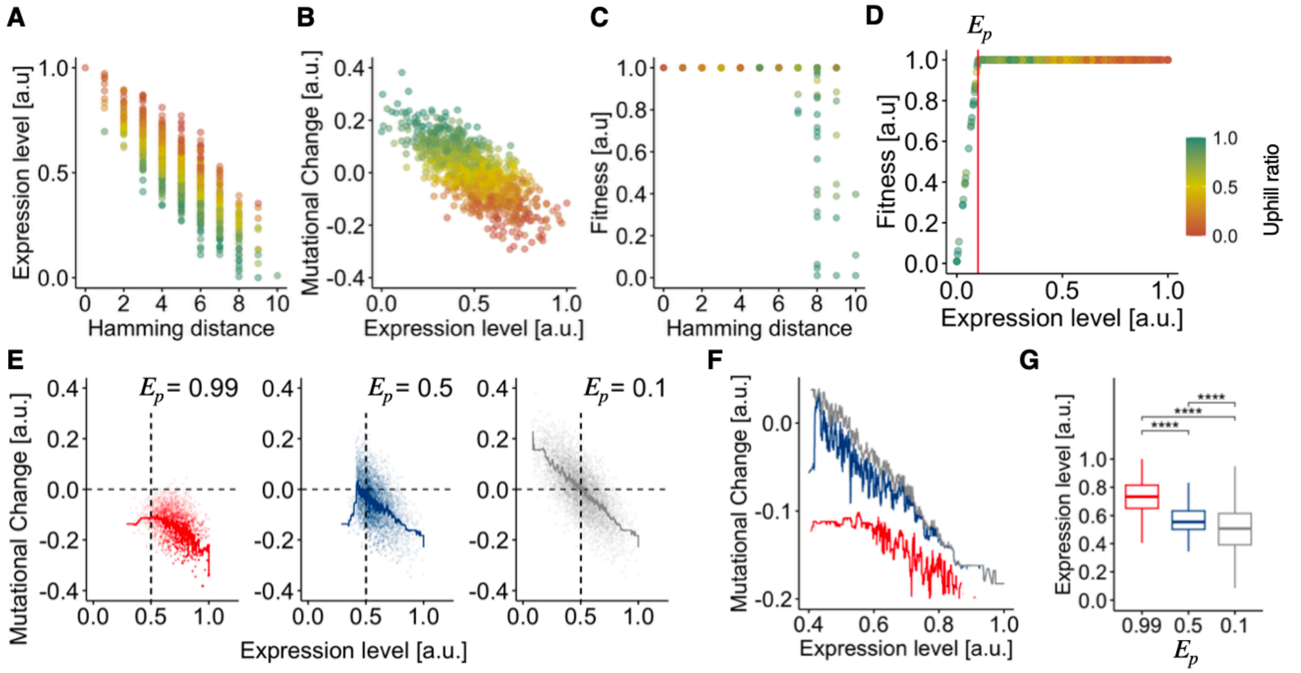


**Fig. S10: Numerical simulation of promoter evolution using the RMF model with linear-plateau relationship between fitness and expression level.**

(**A**, **B**) Representatives of expression landscape (**A**) and mutational changes in expression level (**B**). (**C**) Representative fitness landscape using the linear-plateau model for mapping from expression level to fitness. The linear-plateau model assumes two phases for the relationship between fitness and expression level: a positive linear phase and a flat plateau. The joint point of the two phase was characterized by $E_p$. A smaller $E_p$ corresponds to a wider plateau range over expression levels, while a larger $E_p$ results in a narrower plateau range at higher expression levels. Thus, $E_p$ restricts the range of expression levels with higher fitness. (**D**) Relationship between fitness and expression level in panel (**C**) ($E_p$= 0.1). (**E**) Mutational changes of the evolved isolates obtained through evolutionary simulation using the standard Wright-Fisher model ($E_p$= 0.99, 0.5, and 0.1 from left to right). Solid lines represent the running medians. Dashed lines are guides for the eye. (**F**) Enlarged figure of the running medians in (**E**). (**G**) Expression level of the evolved isolates. Asterisks represent significance levels of the adjusted p-value (the BH method) in the Wilcoxon test (****: $p \leq 0.0001$).

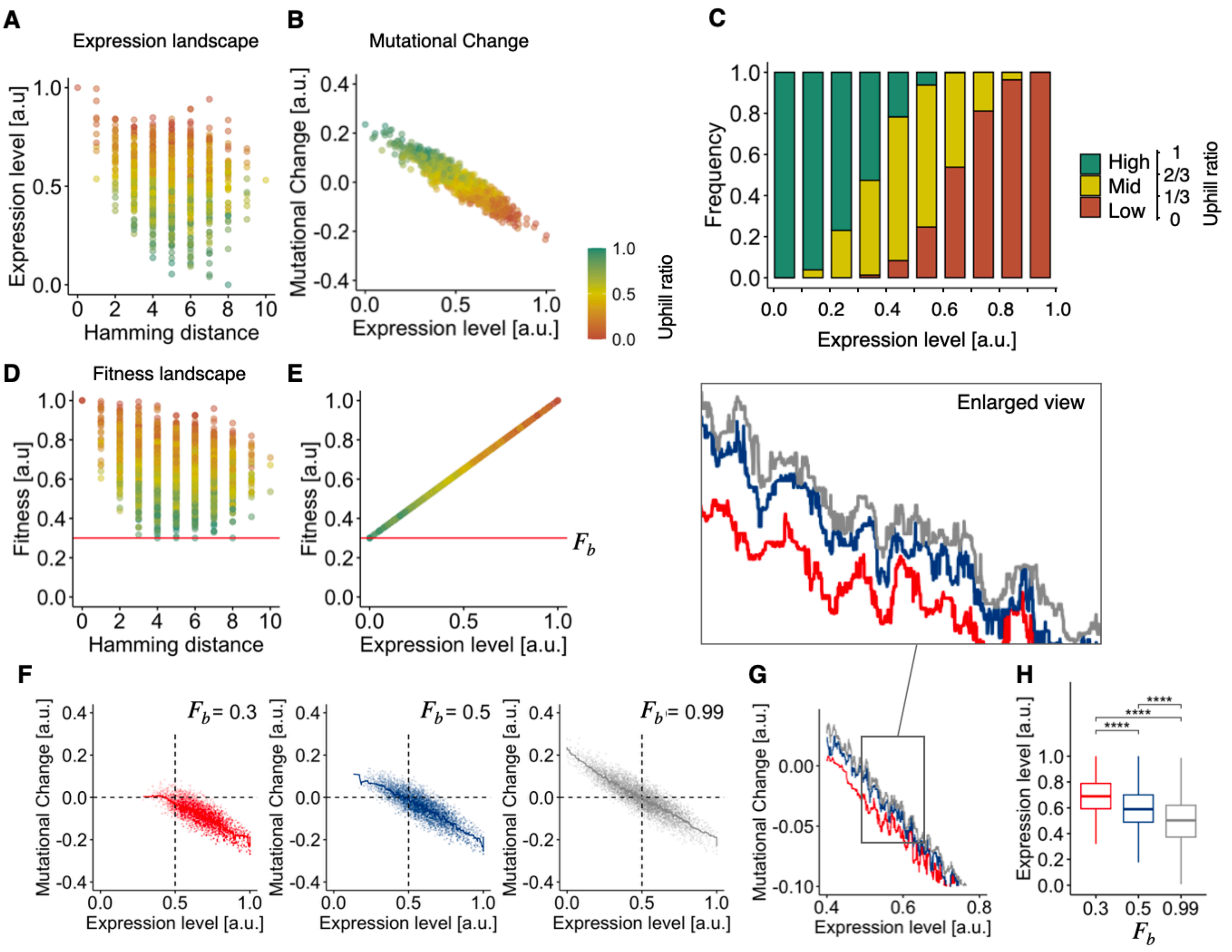


**Fig. S11: Numerical simulation of promoter evolution using the NK model.**

(**A**,**B**) Representative expression landscape (**A**) and mutational changes in expression level (**B**). (**A**) The expression landscape was constructed using the NK model(Kauffman and Weinberger 1989). The Hamming distance represents the distance from the genotype with global optima in terms of expression level and fitness. (**C**) Relationship between the uphill ratio and expression level. Uphill ratio and expression level were divided into three classes (low, mid, and high) and ten classes, respectively in panel **C**. (**D**, **E**) Representatives of fitness landscape (**D**) and relationship between fitness and expression level (**E**) ($F_b$= 0.3, red lines). (**F**) Mutational changes for the isolates obtained through evolutionary simulation using the standard Weight-Fisher model. Ten independent expression/fitness landscapes were generated for each condition of basal fitness ($F_b$=0.3, 0.5, 0.99). In each fitness landscape, 600 independent evolutionary simulations were examined and the initial genotypes were randomly selected. The most abundant mutants within the population after 30 generations were isolated in each run. The solid lines represent running medians. The dashed lines are guides for the eye. (**G**) Enlarged figure of the running medians in (**F**). (**H**) Expression level of the evolved isolates. Asterisks represent significance levels of the adjusted p-value (the BH method) in the Wilcoxon test (****: $p \leq 0.0001$).

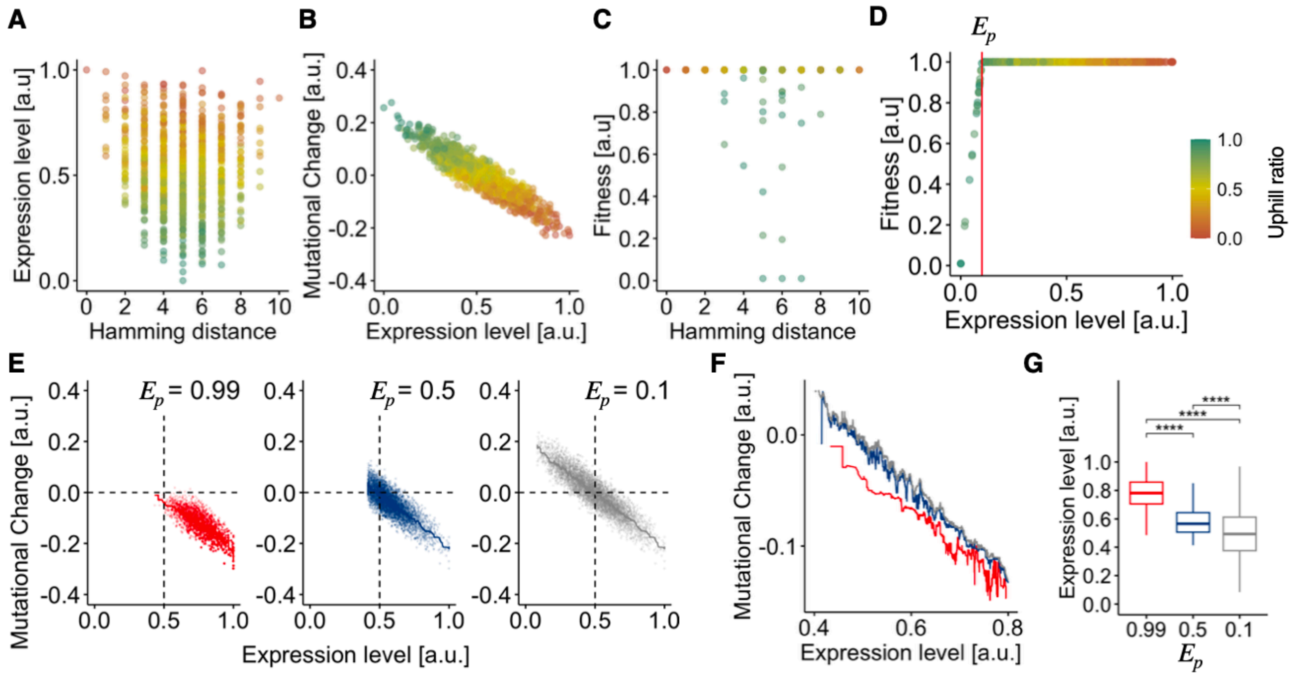


**Fig. S12: Numerical simulation of promoter evolution using the NK model with linear-plateau relationship between fitness and expression level.**

(**A**, **B**) Representatives of expression landscape (**A**) and mutational changes in expression level (**B**). (**C**) Representative fitness landscape using the linear-plateau model for mapping from expression level to fitness ($E_p$= 0.1). (**D**) Relationship between fitness and expression level in panel (**C**). (**E**) Mutational changes of the evolved isolates obtained through evolutionary simulation using the standard Wright-Fisher model ($E_p$= 0.99, 0.5, and 0.1 from left to right). Solid lines represent the running medians. Dashed lines are guides for the eye. (**F**) Enlarged figure of the running medians in (**E**). (**G**) Expression level of the evolved isolates. Asterisks represent significance levels of the adjusted p-value (the BH method) in the Wilcoxon test (****: $p \leq 0.0001$).

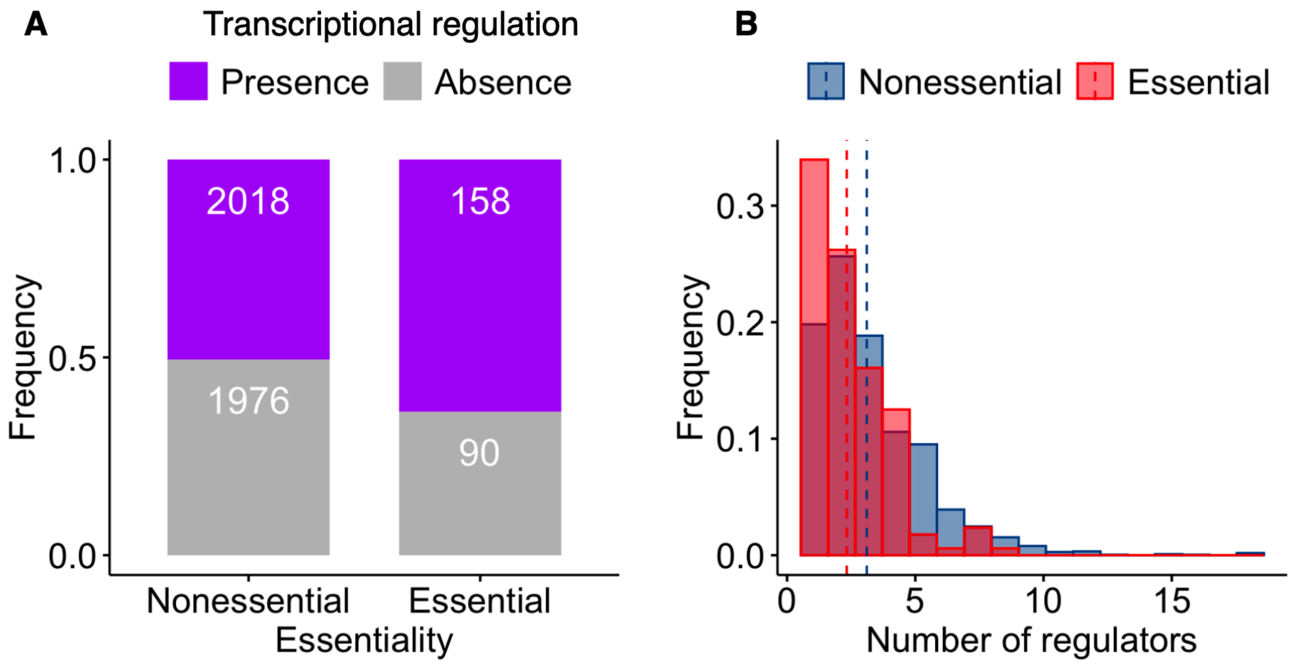


**Fig. S13: Number of transcriptional regulations**

(**A**) Frequency of genes that received no (grey) or more than zero (purple) known transcriptional regulation in each essentiality category. The know regulatory relationship between genes and transcriptional regulators, including all transcription factors and sigma factors, was based on RegulonDB and EcoCyC databases. Essential (248) and nonessential (3994) genes coding for proteins present in the MG1655 strain are shown. (**B**) Distribution of the number of unique regulators by which genes were regulated among the genes that received at least one known transcriptional regulation (purple in pane **A**). The y-axis represents the frequency of each fraction divided by the number of genes that received at least one known regulation (2018 and 158 genes for nonessential and essential, respectively). The red and blue dashed lines represent the mean values.

### II. Supplementary Note

#### Homology of the promoters

We assessed sequence homology among the promoter regions across the three promoter groups (essential, nonessential, and synthetic promoters). A phylogenetic tree was constructed based on a multiple sequence alignment (MSA) of the posterior 73 bp sequences from the 3' ends of the promoters (**supplementary fig. S3B**). External branches of the phylogenetic tree appeared notably long (0.64~) relative to the internal branches, indicating substantial sequence divergence among all pairs of promoters, irrespective of the group. In addition, the maximum size of a clade containing only members from the same group was four at most, suggesting the absence of large solitary clusters specific to any promoter group (**supplementary fig. S3B**). Further confirmation from the blastn(Altschul, et al. 1990) search indicated that synthetic promoters did not align with sequences from existing organisms, supporting their synthetic nature.

While the above MSA provided insights into the diversity of promoter sequences, it offered a coarse-grained analysis based on the overall promoter sequences. Thus, it may potentially underestimate the local similarities in partial sequences responsible for promoter activity. Notably, bacterial natural promoters contain core conserved elements, namely the –35/–10 elements(Gruber and Gross 2003). To explore potential biases at the element level among natural promoters, we focused on core elements recognized by sigma 70, the primary sigma factor in *E. coli*. Using sequences extracted from the RegulonDB database, we constructed a phylogenetic tree based on concatenated –35 and -10 element sequences(Paget and Helmann 2003) (**supplementary fig. S3C**). No significant clustering was observed for essential or nonessential promoters. Additionally, we calculated the Levenshtein distance(Levenshtein 1966) between the concatenated sequences of the two elements and a consensus sequence (TTGACATATAAT)(Shimada, et al. 2014) derived from the –35 ((–35)TTGACA(–30)) and -10 ((–12)TATAAT(–30)) consensus elements (**supplementary fig. S3D**). Statistical analysis revealed no significant difference in this measure between essential and nonessential promoters.

In summary, the three promoter groups exhibited minimal bias in terms of sequence homology, which is conducive for detecting potential signatures of natural selection for essentiality from their mutational effects.
